# Breathing modulates network activity in frontal brain regions during anxiety

**DOI:** 10.1101/2024.06.21.600015

**Authors:** Ana L.A. Dias, Davi Drieskens, Joseph A. Belo, Elis H. Duarte, Diego A. Laplagne, Adriano B.L. Tort

## Abstract

Anxiety elicits various physiological responses, including changes in respiratory rate and neuronal activity within specific brain regions such as the medial prefrontal cortex (mPFC). Previous research suggests that the olfactory bulb (OB) modulates the mPFC through respiration-coupled neuronal oscillations (RCOs), which have been linked to fear-related freezing behavior. Nevertheless, the impact of breathing on frontal brain networks during other negative emotional responses, such as anxiety-related states characterized by higher breathing rates, remains unclear. To address this, we subjected rats to the elevated plus maze (EPM) paradigm while simultaneously recording respiration and local field potentials in the OB and mPFC. Our findings demonstrate distinct respiratory patterns during EPM exploration: slower breathing frequencies prevailed in the closed arms, whereas faster frequencies were observed in the open arms, independent of locomotor activity, indicating that anxiety-like states are associated with increased respiratory rates. Additionally, we identified RCOs at different frequencies, mirroring the bimodal distribution of respiratory frequencies. RCOs exhibited higher power during open arm exploration, when they showed greater coherence with breathing at faster frequencies. Furthermore, we confirmed that nasal respiration drives RCOs in frontal brain regions, and found a stronger effect during faster breathing. Interestingly, we observed that the frequency of prefrontal gamma oscillations modulated by respiration increased with heightened anxiety levels and breathing frequency. Overall, our study provides evidence for a significant influence of breathing on prefrontal cortex networks during anxious states, shedding light on the complex interplay between respiratory physiology and emotional processing.

**Significance Statement:** Understanding how breathing influences brain activity during anxious states could pave the way for novel therapeutic interventions targeting respiratory control to alleviate anxiety symptoms. Our study uncovers a crucial link between respiratory patterns and anxiety-related neural activity in the brain. By investigating the interplay between breathing, neuronal oscillations, and emotional states, we reveal that anxiety induces distinct respiratory patterns, with higher breathing rates correlating with anxious behavior. Importantly, we demonstrate that respiration drives oscillatory activity in the prefrontal cortex, and this effect is potentiated during the fast breathing associated with anxiety. Furthermore, the breathing cycle modulates the emergence of prefrontal gamma oscillations differentially across anxiety levels. This discovery sheds new light on the intricate relationship between respiratory physiology and emotional processing.

## Introduction

Neuronal oscillatory activity can be measured extracellularly and linked to various behavioral and cognitive states (Buzsáki and Draguhn, 2004). While brain oscillations typically arise from intrinsic network mechanisms (Goutagny et al., 2009), there is now substantial evidence also showing their generation by extrinsic factors, such as respiration (Tort et al., 2018a; Folschweiller and Sauer, 2021; Sheriff et al., 2021; Heck et al., 2022). The so-called respiration-coupled oscillations (RCOs) are identified through simultaneous recordings of brain and respiratory activity, where the local field potential (LFP) exhibits a power peak and coherence with the recorded respiration signal at the same frequency as the breathing rate (Tort et al., 2018a, 2018b).

The emergence of RCOs is attributed to the airflow through the nasal cavity, which stimulates sensory neurons in the olfactory epithelium capable of detecting both odor and mechanical stimuli (Grosmaitre et al., 2007). Subsequently, these neurons project to the glomerular layer of the olfactory bulb (OB), which is responsible for propagating respiration-paced neuronal activity through olfactory pathways to distant cortical and subcortical brain regions (Nagayama et al., 2014). RCOs are not exclusively involved in olfactory processing nor only present in olfactory areas (Tort et al., 2018a); in fact, RCOs may influence cognitive functions, including motor, perceptual and memory processes (Heck et al., 2022; Allen et al., 2023; Brændholt et al., 2023) and are most prominent in frontal brain regions (Biskamp et al., 2017; Tort et al., 2018b). Furthermore, RCOs at ∼4 Hz frequency have been linked to negative emotional responses, such as freezing behavior (Karalis et al., 2016; Moberly et al., 2018; Bagur et al., 2021; Karalis and Sirota, 2022). The role of RCOs as a modulator of cognitive and emotional processes underscores the influence of breathing over the activity of different brain networks. Of note, respiration modulates the amplitude of gamma (70–120 Hz) in frontal regions (Cenier et al., 2009; Zhong et al., 2017; Cavelli et al., 2020; Sheriff et al., 2021; González et al., 2023a), indicating an influence of breathing cycles over the processing of multimodal information in association areas (Biskamp et al., 2017).

Despite being a global rhythm, RCOs are often overlooked due to the absence of concomitant respiratory recordings in most electrophysiological studies (Tort et al., 2018a). Another challenge in identifying RCOs is the fact that these oscillations have variable peak frequency, which follows the instantaneous breathing rate. Thus, RCOs may be present in the same frequency range as other common oscillations as delta (∼0.5–4 Hz) and theta (∼5–12 Hz) and be confounded with them (Tort et al., 2018b, 2018a). In particular, RCOs might have been overlooked in studies involving fearful and anxiety-like behaviors in rodents that identified oscillations in the delta and theta frequency range (Adhikari et al., 2010; Lesting et al., 2011; Stujenske et al., 2014; Mooziri et al., 2024).

Therapies based on breathing control can alleviate anxiety symptoms in humans (Hofmann and Gómez, 2017), supporting the idea that anxiety relates to alterations in respiratory patterns (Masaoka and Homma, 1997). This relationship is evident through fluctuations in respiratory and heart rates observed in mice subjected to an anxiety-like behavior test (Okonogi et al., 2018). In rodents, fear and anxiety can be distinguished by specific defensive responses, such as freezing and risk assessment, and their respiratory patterns also differ, with anxiety associated with faster breathing (Carnevali et al., 2013; Okonogi et al., 2018) than fear (Moberly et al., 2018; Bagur et al., 2021). Despite these behavioral differences, negative emotional responses, including fear, anxiety and stress, share common neuronal circuits. Namely, the coordinated activity of the amygdala, ventral hippocampus and prefrontal cortex has been related to the expression of fear and anxiety (for a review, see Calhoon and Tye, 2015). In addition, recent findings also implicate the OB in the fear circuitry given that its inactivation has been shown to both abolish RCOs and reduce freezing (Bagur et al., 2021). Since respiration-entrained signals from the OB have been involved in the modulation of fear expression through organizing mPFC activity, respiration is likely to also influence anxiety-like behaviors that rely on similar neuronal circuits.

Based on that, we hypothesized that neuronal oscillations could appear in the frontal brain not only coupled with the slow breathing characteristic of freezing behavior but also with the faster respiratory rhythms observed during anxiety-like behavior. To test this, we conducted combined behavioral, respiratory, and LFP recordings in Wistar rats subjected to the elevated plus-maze (EPM) task, a common paradigm to evoke anxiety-like behavior (Walf and Frye, 2007). We found that respiratory activity tracks anxiety-like behavior within the EPM task and moreover drives RCOs and modulates gamma oscillations in frontal regions depending on anxiety levels. Our results provide evidence that respiration in the theta frequency range influences the activity of prefrontal cortex networks during anxious states.

## Materials and methods

### Ethics statement

All procedures were performed in accordance with standard ethical guidelines and were previously approved by the Ethics Committee for Animal Experimentation of the Federal University of Rio Grande do Norte under protocol number 062.070/2017.

### Animals

Male Wistar rats (3 months old, n = 7), from the animal facility of the Brain Institute of the Federal University of Natal, were individually housed with controlled temperature (24 ± 1°C) under a 12 light-dark cycle. The experiments were conducted at the beginning of the dark phase. The animals were kept in polyethylene cages with free access to water and food. After the completion of the recordings (see below), the animals were euthanized using an overdose of isoflurane for brain perfusion. All efforts were made to minimize the pain, suffering, or discomfort of the animals.

### Behavioral apparatus

For the elevated plus maze (EPM) task, we utilized an apparatus in “+” shape elevated above the ground with a central area and four arms (EPM: 112 cm length x 12 cm width). Two of these arms had black walls (40 cm high) arranged perpendicular to the other two arms (Figure 1A). In each experimental session, the animal was placed in the central area and allowed to explore the apparatus freely for 10 minutes. Animals were subjected to two EPM task sessions separated by 24 hours. At the end of each session, the floor and walls of the apparatus were cleaned with a 5% alcohol solution. The task was conducted in a dim room with a camera (Point Grey) positioned above the maze recording at 30 frames per second.

**Figure 1.**
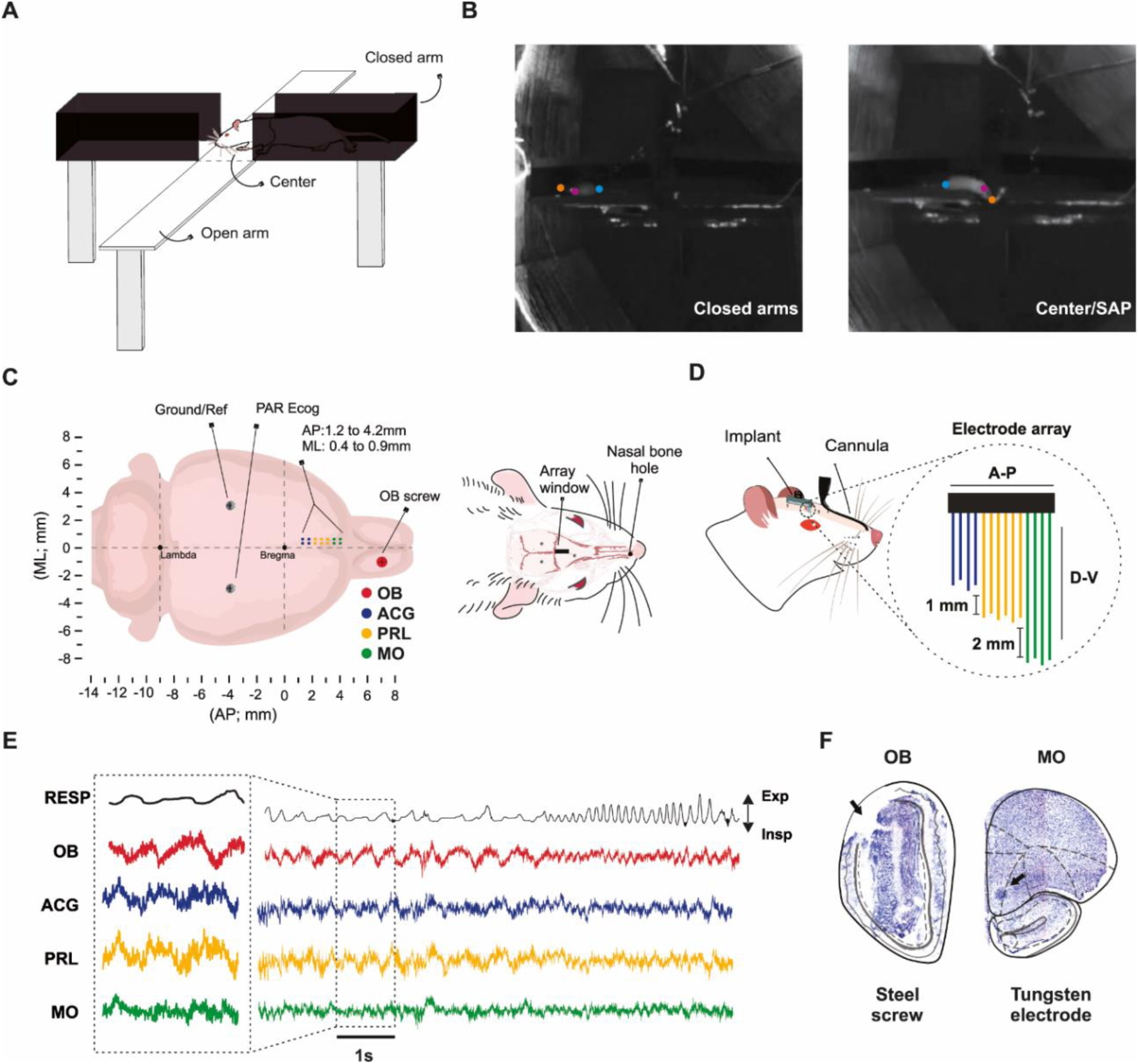
Experimental setup. **(A)** Schematic view of the elevated plus maze (EPM) test. **(B)** An example of DeepLabCut (DLC) tracking when the animal was in the closed arm (left) or performing a stretch-attend posture (SAP) in the center of the EPM (right). **(C)** Schematic illustration of the recording configuration. ACG: anterior cingulate; AP: anteroposterior; Ecog: electrocorticogram; ML: mediolateral; MO: medial orbital; OB: olfactory bulb; PRL: prelimbic; Ref: reference. **(D)** Illustration of the electrode array and of the pressure cannula to record the respiratory signal. **(E)** Example of respiration (RESP) and local field potential (LFP) recordings. **(F)** Histological verification of the recorded areas.

### Surgical procedures

Anesthesia was induced with vaporized isoflurane (5%), followed by intraperitoneal injections of ketamine (100 mg/kg) associated with xylazine (4 mg/kg), and then maintained with vaporized isoflurane (1.5 - 2.5%). After anesthetic induction, the animals were shaved, aseptically prepared and fixed with ear-bars in the stereotaxic apparatus (RWD Life Science); a scalp incision was then made to expose the skull. To record multiple brain areas, the animals were implanted with custom-designed electrode arrays with 14 tungsten wires (75 μm in diameter, California Fine Wire Co, USA). The positioning and arrangement of the array (2×7; inter-electrode distance: 500 μm, see Figure 1C) allowed us to record the electrical activity from the medial prefrontal cortex (AP: 1.2 - 4.2 mm; ML: 0.4 - 0.9 mm; apart) (Paxinos & Watson, 2007), including the medial orbital (MO), prelimbic (PRL), and anterior cingulate (ACG) regions (Figure 1F). We also recorded an ECoG signal from the right olfactory bulb with a stainless-steel screw resting over its dorsal surface (OB; AP: 7.5 mm, ML: -1 mm) and used a second screw over the left parietal cortex as reference (PAC; AP: -4 mm, ML: +3 mm). All electrode signals were acquired at 12207 Hz with a PZ2 PreAmp connected to an RZ2 Bioamp (Tucker Davis Technologies, USA)

To record respiratory activity, we implanted a stainless steel cannula syringe (18 gauge blunted needle, with the tip removed, BD USA) into the right nasal cavity. To ensure optimal positioning without obstructing the animal vision, we mechanically twisted the cannula into an “S” shape (Figure 1D). Subsequently, we drilled a hole into the nasal bone (approximately 1.5 cm posterior to the nasal vestibule) using a surgical drill. During recordings, we reversibly connected the other end of the cannula to a head-mounted pressure sensor (24PCAFA6G, Honeywell, Morristown, USA) amplified with an AD627A amplifier circuit (Analog Devices, USA). We digitized the pressure signal at 12207 Hz with the RZ2 system.

### Perfusion and histology

After completing the recordings, an analysis of the histological sections was carried out to confirm electrode positioning. To that end, the brains were removed, stored and then cut and mounted on slides of glass. Sections were viewed and photographed under a microscope equipped with fluorescence (Zeiss).

### Data analysis

All analyses were performed using MATLAB (Mathworks), unless otherwise stated.

#### Video Tracking

We tracked the *x* and *y* coordinates of the animals’ head positions within the EPM from the overhead videos using DeepLabCut (DLC) (Nath et al., 2019).

#### Behavior Analysis

Based on the *x* and *y* coordinates obtained by the DLC, custom-written Matlab routines were used to quantify the animal speed and distance traveled, as well as to classify the animal location into 5 maze regions: open arms (up and down), closed arms (right and left), and center area.

To calculate the total time spent in each maze region, we multiplied the number of frames in each position by 1/(30 fps). For the distance traveled, we first converted pixels to centimeters using a factor estimated from the length of the arms (112 cm) and the associated number of pixels and then computed the Euclidean distance between consecutive frames. The instantaneous animal speed was obtained by multiplying the distance by 30 fps. Positional coordinates associated with speed values exceeding 100 cm/s, which represented only 0.1 to 2% of each animal data, were manually “deleted” (converted to “NaN”) as these were artifacts resulting from the loss of tracking by DLC. After this removal, we applied the Matlab ’smooth’ function to interpolate NaN values using neighboring valid data points, thereby ensuring a smoother representation of positional coordinates. Both traveled distance and speed were recomputed after smoothing the positional coordinates. Finally, instantaneous speed values were averaged using 1-s windows to allow comparisons with the respiratory frequency (which was estimated using 1-s windows; see below and Figure S3).

To assess anxiety-like behavior, we analyzed the time spent in the center area & open arms compared to the closed arms (Carobrez and Bertoglio, 2005). In addition, we also analyzed risk assessment behavior by quantifying the frequency of stretch-attend postures (SAP). These were identified by visual inspection of video recordings and defined to occur when the animal was in the center area stretching its body, extending its head and forepaws into the open arms while the rest of its body remained in the center or closed arms (see Figure 1B right). Since animals performed SAP while in the center area, we combined the time spent in the center and open arms to define the total time spent in an anxiety zone.

#### Power Spectrum Density

The LFP and ECoG signals were low-pass filtered at < 500 Hz and, along with the respiration signal, subsampled using a factor of 10 (new sampling rate: 1220.7 Hz). Prior to computing power spectrum densities (PSDs), we performed a piecewise detrending of the electrophysiological and respiration signals using non-overlapping 1-second windows (i.e., each 1-s window was independently detrended). Then, for each time window a PSD was computed using a Hamming window of the same length (1 s) through the *pwelch.m* function. The PSD of the individual windows were stored in a matrix, thus giving rise to a spectrogram. To reduce variability among channels and animals, each spectrogram was normalized by dividing the PSD values by a normalization factor defined as the average power below 12 Hz (all task period considered). The spectrogram of each brain region was taken as the average normalized spectrogram over all channels in the region. To compute the PSDs according to maze location, for each session and brain region we averaged all spectrogram columns (i.e., all PSDs) whose timestamp was associated with that location (e.g., permanence in the closed arms).

#### Respiratory Frequency

To estimate the instantaneous respiratory frequency, we first computed a spectrogram of the respiratory signal using non-overlapping 1-second windows as described above. Then, for each time window, the power distribution was normalized by its maximum power value. The respiratory frequency of each temporal window was then defined as the frequency associated with the highest power value (in this case, 1 after normalization; see Figure 3A).

#### Phase Coherence

To calculate phase coherence of the LFP signal with breathing, we used the *mschore.m* function (1 second windows with 50% of overlap). For coherence analysis taking into account the location of the animal in the EPM (closed arms vs. center + open arms), a concatenation was first performed of the LFP and the respiratory signals using the timestamps of interest. Similar to the power analysis, the phase coherence of each brain region was taken as the average coherence spectrum over all channels in the region.

To compare the coherence between the LFP and breathing against chance, surrogate analyses were employed. For this, surrogate values were generated by computing coherence spectra between the respiratory signal and the LFP temporally shifted in a circular manner; we employed 20 surrogates for each analyzed LFP. For each surrogate, the magnitude of the shift was randomly determined between 1 and 2 seconds, following a uniform distribution. For each region, the actual mean phase coherence was compared with the distribution of surrogate values in the corresponding frequency range.

#### Phase-Amplitude Coupling

To calculate phase-amplitude coupling, we used the method described by Tort et al. (2010). To that end, we filtered the LFP to obtain the faster frequency components (20 – 200 Hz with a step of 5 Hz; bandwidth = 20 Hz) and the respiration signal to obtain the slower frequency components (0 – 12 Hz with a step of 0.5 Hz; bandwidth = 2 Hz). Filtering was performed using the *eegfilt.m* function from the EEGLAB toolbox (Delorme and Makeig, 2004). The phase and amplitude components were then estimated based on the Hilbert transform using the *hilbert.m* function. The slow component phase was binned into 18 bins of 20° degrees. The comodulograms were obtained by computing the entropy-based modulation index (MI) for each slow-fast frequency pair using all timestamps of interest (i.e., all time periods associated with a given EPM region). Before computing the average, each comodulogram was z-scored to reduce variability among animals.

#### Granger Causality

Granger causality analysis was conducted using the Multivariate Granger Causality MATLAB Toolbox (Barnett and Seth, 2014). The LFP and respiration signals were further downsampled by a factor of 10 (new sampling rate = 122 Hz). Additionally, a piece-wise detrending procedure was applied in the data (similarly to the power spectrum estimation). The variational autoregressive model order was set to 40.

#### Statistical Analysis

Group data was represented as mean ± SEM. Statistical differences were assessed through paired t-tests using *ttest.m*, and two-way or repeated measures ANOVA using *anovan.m*. Values with p < 0.05 were considered statistically significant.

## Results

In this study, we sought to examine the relationship between brain activity and nasal breathing during anxiety-like behavior. To that end, we performed local field potential recordings from the olfactory bulb (OB) and medial prefrontal cortex (mPFC) regions of Wistar rats exposed to the elevated plus maze (EPM) test while tracking their respiratory activity through an intranasal pressure cannula (Figure 1).

### Higher exploration of open arms during the first minutes of the EPM task

To study anxiety-like behavior, we employed the EPM test, which is based on the innate tendency of the rodents to avoid open elevated areas (i.e., the open arms of the maze) while contrasting with their inherent curiosity to explore new environments. In this task, the animals feel safer when they are in the closed arms, while open arm exploration can be taken as a risk-assessment behavior associated with higher anxiety (Pellow et al., 1985; Lister, 1987). Thus, in this work we fragment the analysis of EPM exploration depending on animal location into closed arms vs. open arms + center area. Notice that this “within-session” approach, also used in previous electrophysiological studies (Adhikari et al., 2010; Mooziri et al., 2024), contrasts with classical EPM analyses that take a single metric (e.g., number of open arm entries) for the whole session.

During the 10-minute exposure to the maze, animals exhibited a higher tendency to explore the open arms and center area in the first ∼3 minutes of the test compared to the later periods (Figure 2A; repeated measures ANOVA over minute-by-minute time bins, F(9,117) = 8.97, p < 10^-9^, n = 14 sessions). To maximize data yield, each animal (n = 7) was subjected to two EPM task sessions separated by 24 hours. We thus conducted a statistical analysis to assess whether the repeated EPM exposure influenced the anxiety levels as measured by the exploration time of different maze zones. Two-way ANOVA using maze location (center + open vs closed arms) and EPM session number (1 and 2) as factors showed a major effect of location (F(1,24) = 612.3, p < 10^-17^) but no interaction with the experimental session day (F(1,24) = 0.06, p = 0.81). These results show that the time spent in the closed arms is similar between experimental sessions, indicating a consistent manifestation of anxiety-like behavior in the task irrespective of prior maze experience. Similarly to the profile of exploration in the open arms, the time course of the distance traveled in the maze showed higher activity in the first 3 minutes (Figure 2C, F(9,117) = 5.42, p < 10^-5^, repeated measures ANOVA). Also as open arm exploration, the animals did not show differences in the total distance traveled between both experimental sessions (Figure 2C inset, t(6) = -1.67, p = 0.15, paired t-test). Finally, when quantifying risk assessment through the execution of stretch-attend posture (SAP), we also found a profile in which the animals tended to perform more SAP during the first minutes of the task (Figure 2D), with no difference between the two experimental sessions (Figure 2D inset, t(6) = -0.60, p = 0.57, paired t-test). Of note, the SAP behavior typically occurred when the animals were exploring the center of the arena, with their body stretching toward the open arms.

**Figure 2.**
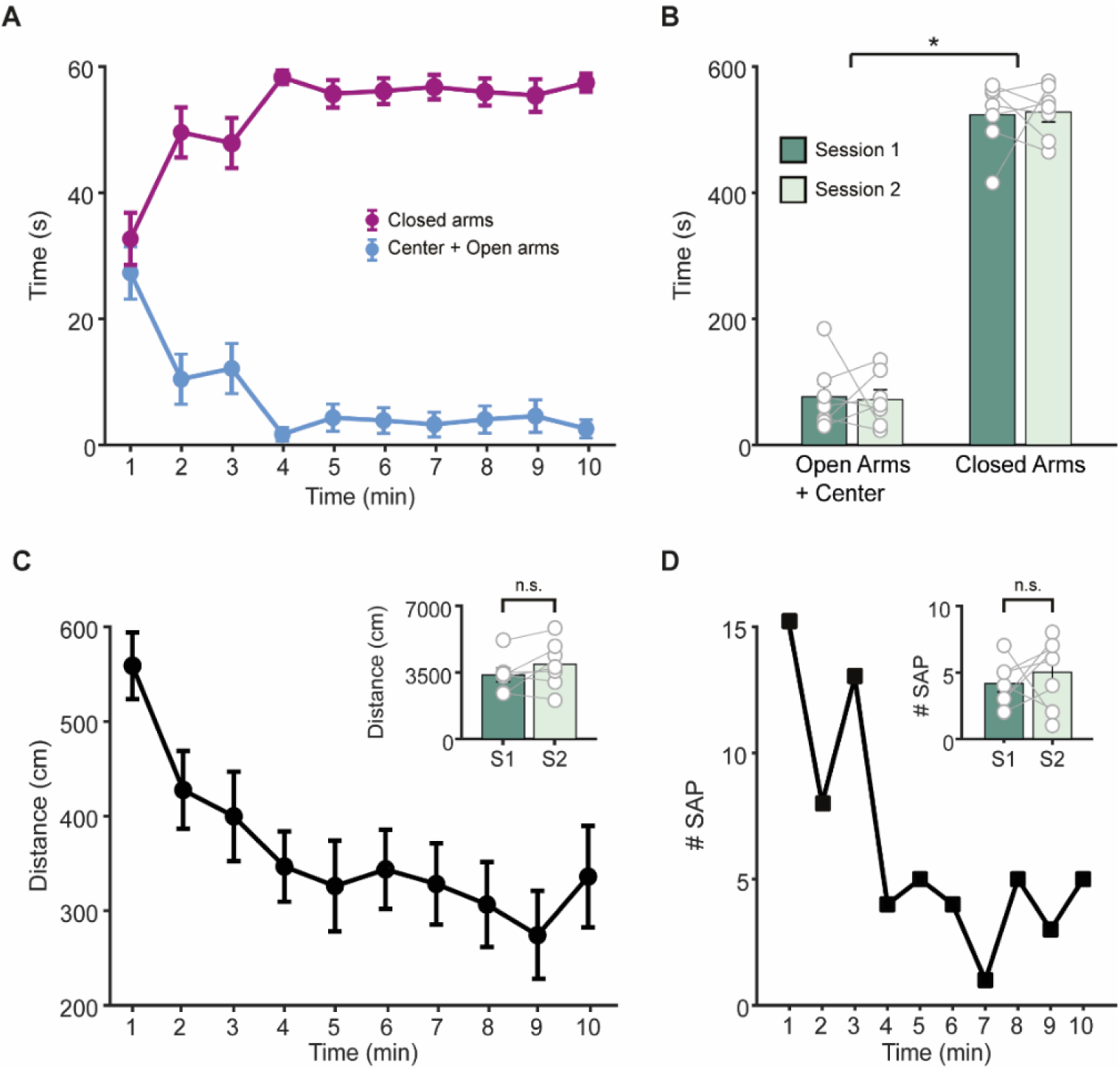
Similar behavior across EPM sessions. **(A)** Minute-by-minute time course of permanence in center + open vs. closed arms (N = 14 sessions from 7 rats). **(B)** Total time spent in center + open vs. closed arms during sessions 1 and 2. *p < 0.0001 for the maze region factor and p = 1 for the session factor, two-way ANOVA. **(C)** Time course of distance traveled in the EPM. Inset shows the total distance traveled in sessions 1 (S1) and 2 (S2). p = 0.15, paired t-test. **(D)** Sum of the number of stretch-attend postures (SAP) per minute across 14 sessions. The inset shows the mean number of SAP episodes in S1 and S2. p = 0.57, paired t-test. n.s., not significant.

In summary, our results show that in the initial 3 minutes of the task animals explore more the open arms & center, exhibit a higher frequency of SAP behavior, and move more compared to the later periods. Besides that, the animals spent more time in the closed arms during the 10 minutes of the task and displayed consistent behaviors across the two sessions in the EPM.

### Behavioral state-dependent respiratory rate in the EPM

After analyzing the general behavioral profile of the animals in the EPM, we moved on to investigate potential relationships between behavior in the maze and the respiratory dynamics tracked through the nasal pressure sensor (Figure 3A top panel). We started by conducting power spectral density (PSD) analyses to examine the main frequency bands present in the respiratory signal. To that end, we computed PSDs in non-overlapping 1-second windows (Figure 3A middle panel). The instantaneous respiratory frequency of each window was defined as the one with the highest power (to aid visualization of the peak frequencies in Figure 3A bottom panel, the power spectrum of each window was normalized by the maximal value).

**Figure 3.**
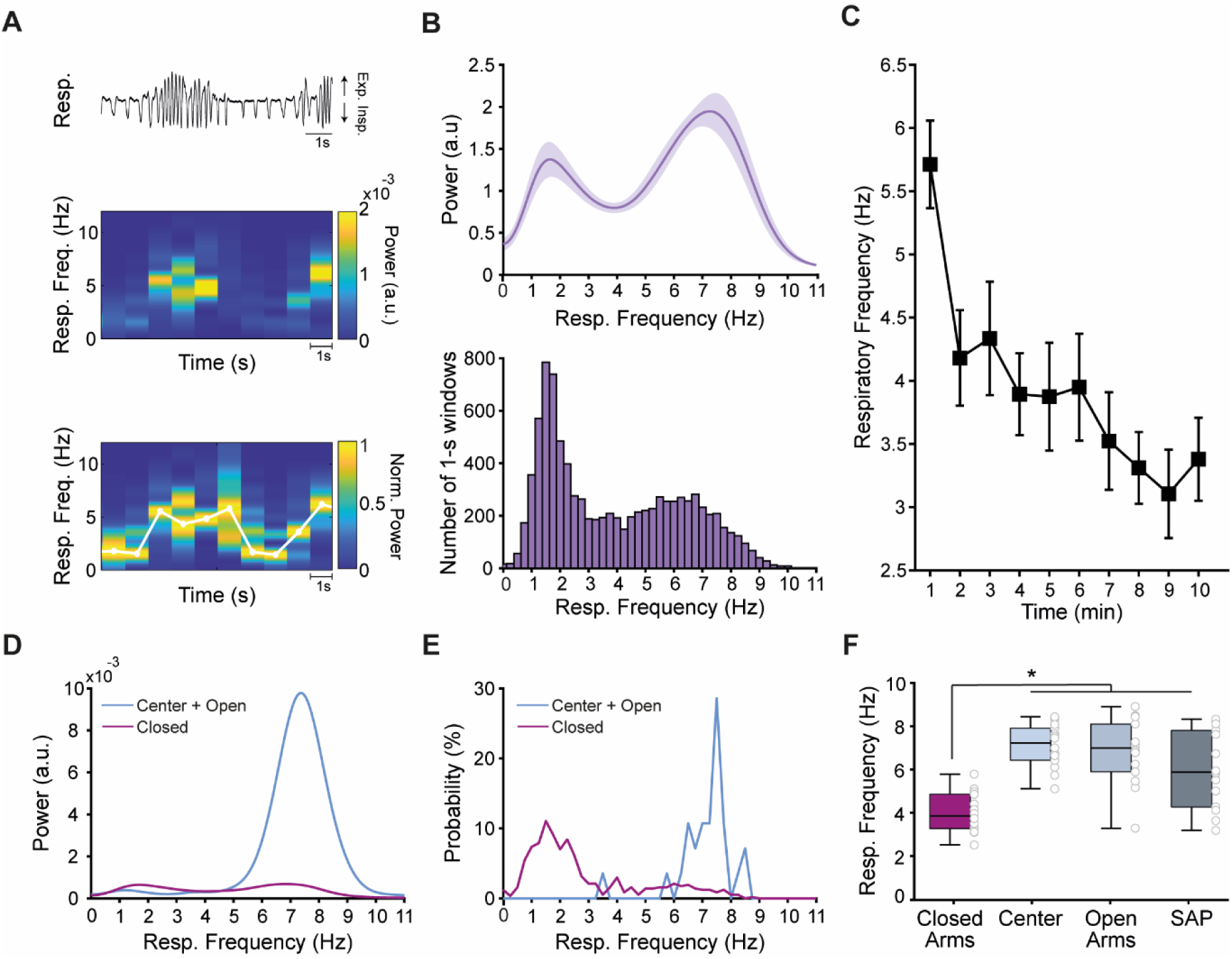
Anxiety-like behavior in the EPM increases the respiratory frequency. **(A)** Representative respiratory signal derived from the nasal air pressure sensor (top) and the corresponding time-frequency power spectrum obtained using non-overlapping 1-s windows (middle). The bottom panel shows the power spectrum of each window normalized by the maximal value. The white line indicates the estimated “instantaneous” respiratory frequency, defined as the peak power frequency in each window. Exp.: expiration; Insp.: Inspiration. **(B)** Power spectrum of the respiratory signal (top; M ± SEM, N = 14 sessions), revealing a bimodal distribution of respiratory rate (see also Figure S1). The histogram (bottom) displays the distribution of the instantaneous respiratory frequencies (pool among sessions). **(C)** Minute-by-minute time-course of respiratory frequency during EPM exposure (M ± SEM, N = 14 sessions). Note the gradual decrease over time. **(D,E)** Power spectrum (D) and probability distribution (E) of respiratory frequency during the exploration of the EPM center + open arms v.s. closed arms in a representative animal. **(F)** Respiratory frequency in the different zones of the maze and during SAP events (N = 14 sessions). Notice increased respiratory rates in the anxiogenic zones of the maze compared to the closed arms. *p < 0.0001, repeated measures ANOVA followed by the Tukey-Kramer test.

When computing the average PSD over the entire task period (i.e., across all time windows), we found that the animals exhibited a bimodal respiratory frequency distribution, with a tendency to breathe either in a slower (∼ 0.5 – 3 Hz) or a faster (∼ 6 – 10 Hz) range (Figure 3B top panel; see also Supplementary Figure S1 for individual animal results). The respiratory power in the faster frequency range was greater than in the slower range; however, during periods of slow breathing, the air pressure signal had lower amplitude, while faster breathing was associated with much higher air pressure (see the example trace in Figure 3A and Figure S1A). Therefore, the higher power observed for the faster than slower breathing does not necessarily mean that animals spent more time breathing at this rate. Thus, to disentangle signal amplitude from its duration, we analyzed the histogram distribution of the peak respiratory frequencies of all time windows. This showed that the animals spent more time breathing at slower frequencies during the EPM task (Figure 3B bottom panel and Figure S1).

When plotting the average respiratory frequency over time, we observed a clear decrease along the 10 minutes of exposure to the EPM (Figure 3C, F(9,117) = 7.09, p < 10^-7^, repeated measures ANOVA). This result was consistent in the two experimental sessions (Figure S2) and resembled the behaviors of exploring the center + open arms and performing SAP, which were also more prevalent in the first minutes of the task (Figure 2A,D). To directly investigate whether changes in respiratory frequency correspond to the different behaviors, we next separately computed the average respiratory frequency when the animals were in different regions of the maze. Interestingly, inspection of both the power spectrum of the respiratory signal as well as of its peak frequency distribution revealed a greater occurrence of slower breathing during the periods in the closed arm, and of faster breathing rates during center + open arm exploration (see Figure 3D and 3E for a representative example). At the group level (Figure 3F), we found a statistically significant difference in respiration rate when the animals were in the closed arms (lower) compared to the open arms, center area and during SAP (F(3,39) = 29.44, p < 10^-9^, repeated measures ANOVA followed by Tukey-Kramer multi comparisons test). Of note, we found a consistent profile across the two sessions in the EPM, wherein there was a significant difference in respiration rate depending on the maze zone (F(1,48) = 13.69, p < 10^-5^) but not between the sessions (F(1,48) = 0.09, p = 0.76) (Figure S2C). These results therefore indicate that the animals breathe at higher frequencies when exhibiting anxious-like behavior, and that the bimodal respiratory rate distribution displayed by the animals relates to maze location.

### Respiratory rate correlates both with locomotor activity and anxiety-like behavior

Upon observing that animals breathe faster during periods in the center + open arms and during SAP, we next investigated whether this result could be associated with putative anxious states or else be entirely explained by changes in locomotor activity. This investigation is motivated by the fact that breathing rate relates to the speed of locomotion (Nguyen Chi et al., 2016).

For locomotion analysis, we computed the average head speed in 1-second temporal windows, that is, the same time windows used to estimate the instantaneous breathing rate (Figure S3A-C). We found that the average speed in the EPM ranged from ∼0.5 to 20 cm/s (Figure S3D). Subsequently, we analyzed the average respiratory frequency as a function of the animal speed and observed a strong positive relationship between these two variables (Figure 4A and Figure S3E), which was stable between the two experimental sessions (Figure S4A).

**Figure 4.**
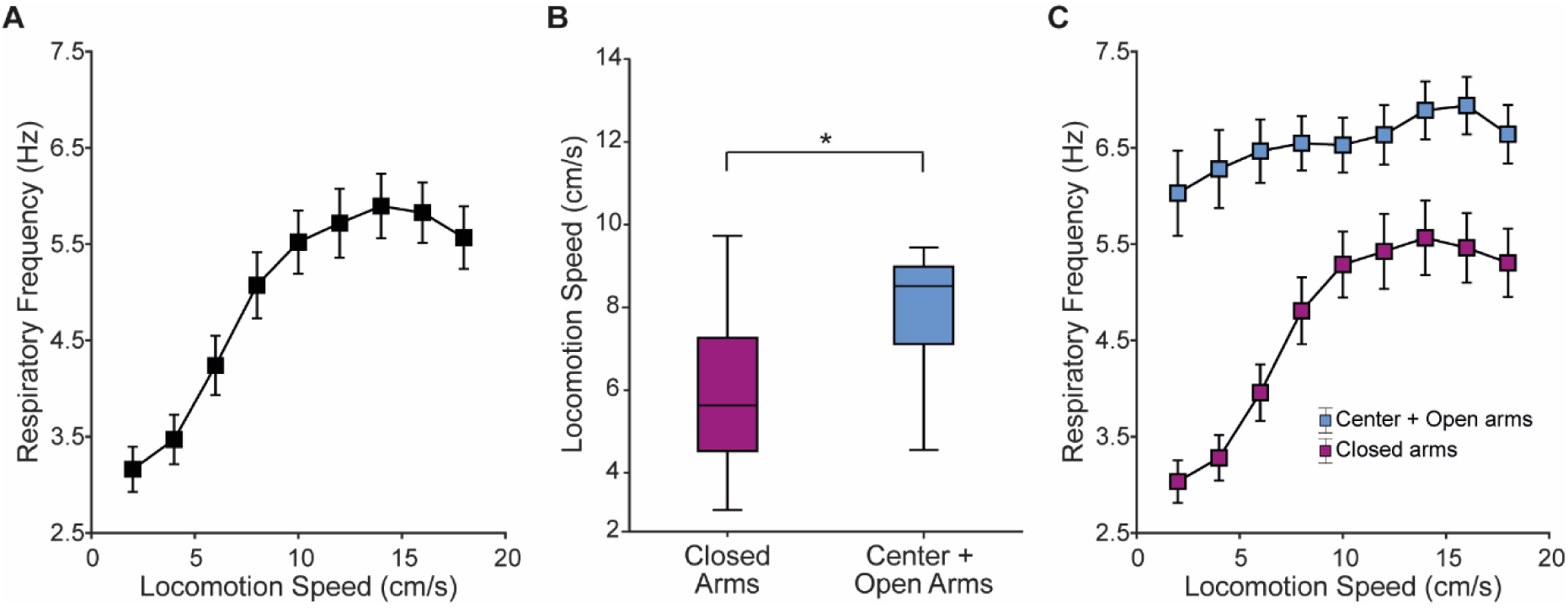
Respiratory frequency predicts putative anxious states irrespective of locomotion speed. **(A)** Mean respiratory frequency (± SEM) as function of the animal locomotion speed in the EPM (N = 14 sessions). **(B)** Average speed during exploration of the EPM center + open arms vs. closed arms (*p <0.01, paired t-test). **(C)** Mean respiratory frequency as a function of the speed plotted separately for when the animal was in the closed arms vs. center + open arms. Note a significant increase in respiratory frequency in the center + open arms independently of the animal’s locomotion speed (p < 0.0001, two-way ANOVA).

Next, we examined the average locomotion speed during periods in the closed arms vs. center + open arms (note that most SAP behaviors occur in the center). Our analysis revealed that animals exhibited higher locomotion speeds in the center + open arms compared to the closed arms (Figure 4B; t(13) = 3.41, p < 0.01, paired t-test). Given this observation, we deemed possible that the increased respiratory frequency during the putative anxious state associated with center + open arm exploration (Figure 3F) was due to differences in speed in the different regions of the maze.

To further explore this possibility, we conducted a two-way ANOVA considering as factors the position in the maze (closed vs. center + open arms) and locomotion speed. We found that both factors influenced the respiration rate (position: F(1,229) = 139.62, p < 10^-24^; speed: F(8,229) = 6.97, p < 10^-7^). Interestingly, when plotting the respiration rate as a function of speed separately for each maze region, we found a major increase when the animals were in the more anxiogenic zones of the maze regardless of the locomotion speed (Figure 4C). For instance, for the lowest speed bin (∼ 2 cm/s), there was a difference of 3 Hz in respiration rate when comparing open vs. closed arm exploration, and the difference in respiration rate was > 1 Hz even at the fastest speeds (Figure 4C). Consistently, the two-way ANOVA employed above also confirmed an interaction between maze position and speed (F(8,229) = 2.52, p = 0.01), which was characterized by a steep increase in respiration rate as a function of speed when the animals wandered in the closed arms, and a lesser effect of speed over locomotion when the animals were in the center + open arms (Figure 4C). Quantitatively, respiration rate increased by ∼66% from the slowest to the fastest speed in the closed arms (from ∼ 3 to ∼ 5 Hz) while it increased by only 8% in the center + open arms for the same speed range (from ∼ 6 to ∼ 6.5 Hz). In all, this result demonstrates that the increased respiratory frequency during center + open arm exploration relates more to anxiety than locomotion speed.

### Respiration-coupled oscillations at distinct frequency bands in the frontal brain

Cerebral rhythmic activity is often categorized by frequency ranges (Buzsáki and Draguhn, 2004). However, recent work has shown that breathing is capable of impinging a neuronal rhythm that defies the classical range definition due to its variable frequency. Namely, the so-called respiration-coupled oscillations (RCOs) have been shown to follow breathing rate, spanning from delta (0.5 – 4 Hz) and theta (4 – 12 Hz) up to low beta (12 – 18 Hz) frequency ranges (Rojas-Líbano et al., 2014; Tort et al., 2018a). Operationally, to identify RCOs, it is necessary (1) for the spectral activity of the local field potential (LFP) to exhibit a power peak in the same frequency as respiration along with (2) phase coupling (synchrony) with the respiratory signal at this same frequency (Tort et al., 2018b, 2018a). Thus, to investigate the presence of RCOs during EPM exploration, we analyzed the LFP power spectrum of the olfactory bulb (OB), prelimbic cortex (PRL), anterior cingulate gyrus (ACG), and medial orbitofrontal cortex (MO), and also measured the LFP phase coherence to the respiratory activity.

Upon analyzing the whole task period (10 minutes), the electrophysiological and respiratory signals exhibited power peaks at the same frequency ranges and also phase coherence levels above chance (as determined through surrogate analysis, see Materials and methods) (Figure 5A and Figure S5). Therefore, based on the operational definition outlined above, these results demonstrate the presence of RCOs in frontal brain regions during exploration of the EPM. Notably, similarly to respiratory activity during the task, the LFP power peaks could be detected within two distinct frequency bands associated with slow (∼0.5 – 4 Hz) and fast (∼5 – 10 Hz) breathing. Of note, as opposed to the respiration power spectrum, the LFP power peak was much higher in the slow than in the fast frequency range, suggesting that the LFP RCOs have larger amplitude during slow breathing, consistent with previous reports (Girin et al., 2021).

**Figure 5.**
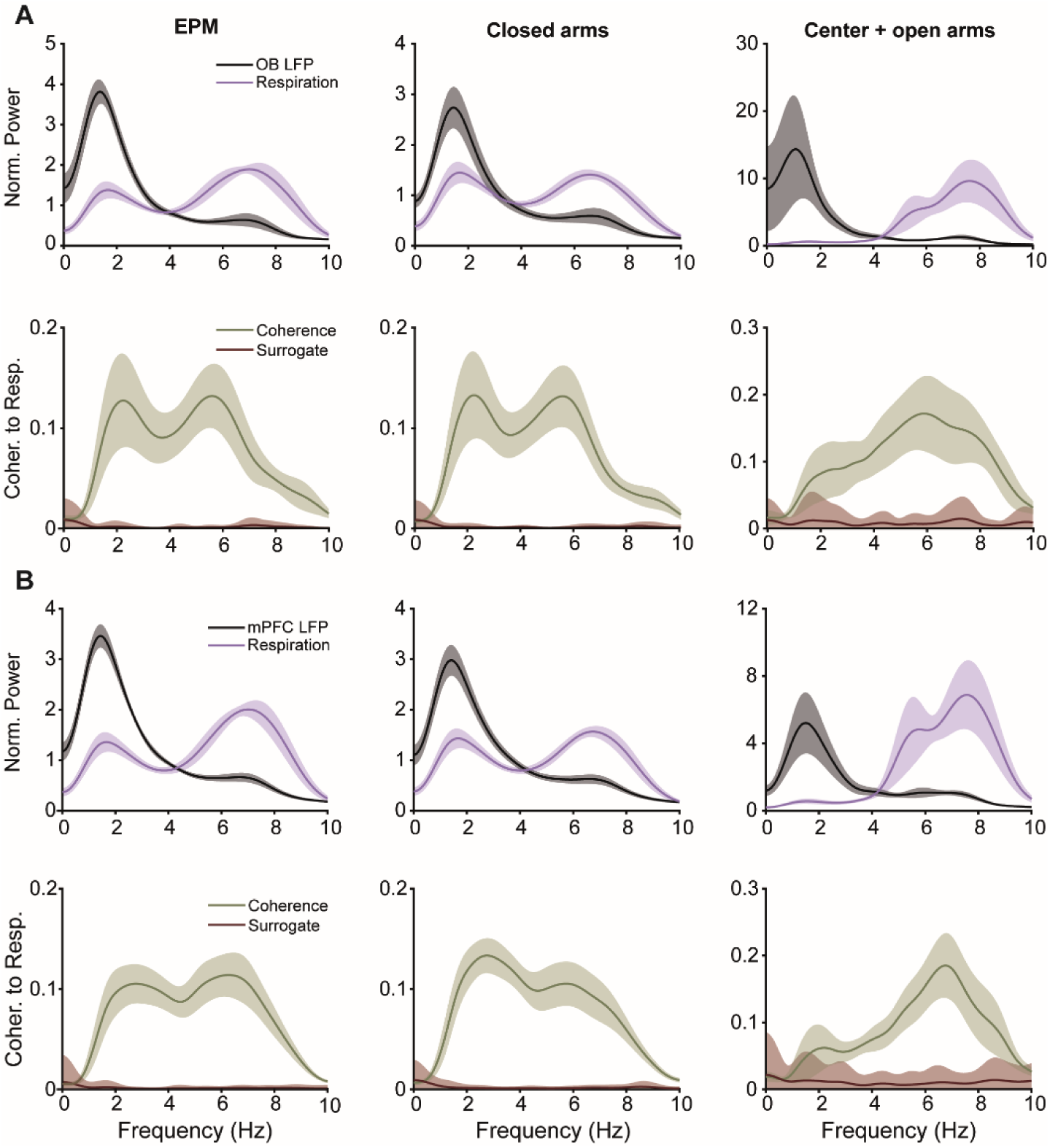
Respiration-coupled LFP oscillations occur at both slow and fast frequencies in the EPM. **(A)** Top: Mean power spectrum density of respiration (purple) and OB LFP (black) computed for the whole task period (left) and for periods when the animal was in the closed arms (middle) or in the center + open arms (right). Bottom: Respiration – OB LFP coherence (green) for the same periods as above. The red trace shows the mean coherence chance levels. **(B)** Same as in A but for the MO LFP (see Figure S5 for the other mPFC regions). The shaded areas indicate ± SEM for the actual spectra and ± 2*SD for the surrogate spectra. Note that the levels of LFP – Resp coherence are much above chance, which defines the presence of respiration-coupled oscillations. Notice further the existence of two peaks corresponding to slow and fast breathing regimes.

Interestingly, when performing spectral and coherence analyses according to animal location in the EPM, LFP power peaks were observed in both frequency bands when the animals were in the closed arms and in the center + open arms (Figure 5). Nevertheless, when the animals explored the center + open arms, coherence to respiration was more pronounced for LFPs within the faster frequency range (Figure 5).

### Higher LFP power at fast respiratory rates during anxious states

To investigate whether the effect of respiration over frontal brain activity is modulated by anxiety-like states, we compared the average LFP power in the closed arms versus the center + open arms. Given that the animals breathe at faster frequencies in the more anxiety-inducing zones (center + open arms), for these analyses we selected only the periods in which the animals breathed at rates exceeding 4 Hz in either maze location. This approach allows for a proper power comparison by excluding periods of slow breathing in the closed arms that are absent in the open arms, thus avoiding dilution of power density values for fast frequencies in the closed arms.

Interestingly, when considering only the periods of faster breathing rates (>4 Hz), the respiratory signal exhibited significantly larger power in the center + open arms compared to the closed arms (Figure 6A, t(11) = 3.88, p < 0.01, paired t-test), indicating that the animals take deeper breaths during anxious states. Noteworthy, while the coherence analyses showed similar phase-locking between the LFP and the respiratory signal during both the closed and center + open arm explorations (Figure 6B; t(8) = 0.55, p = 0.59), the power spectrum revealed a statistically significant higher RCO power at fast frequencies in the anxiety-inducing zone (Figure 6C,D; OB: t(8) = 2.58, p = 0.03; MO: t(9) = 2.40, p = 0.04; PRL: t(11) = 2.92, p = 0.01; ACG: t(11) = 3.52, p < 0.01). Therefore, our results show a greater influence of respiration on prefrontal cortex activity during anxiety states, as reflected by increased RCO amplitude.

**Figure 6.**
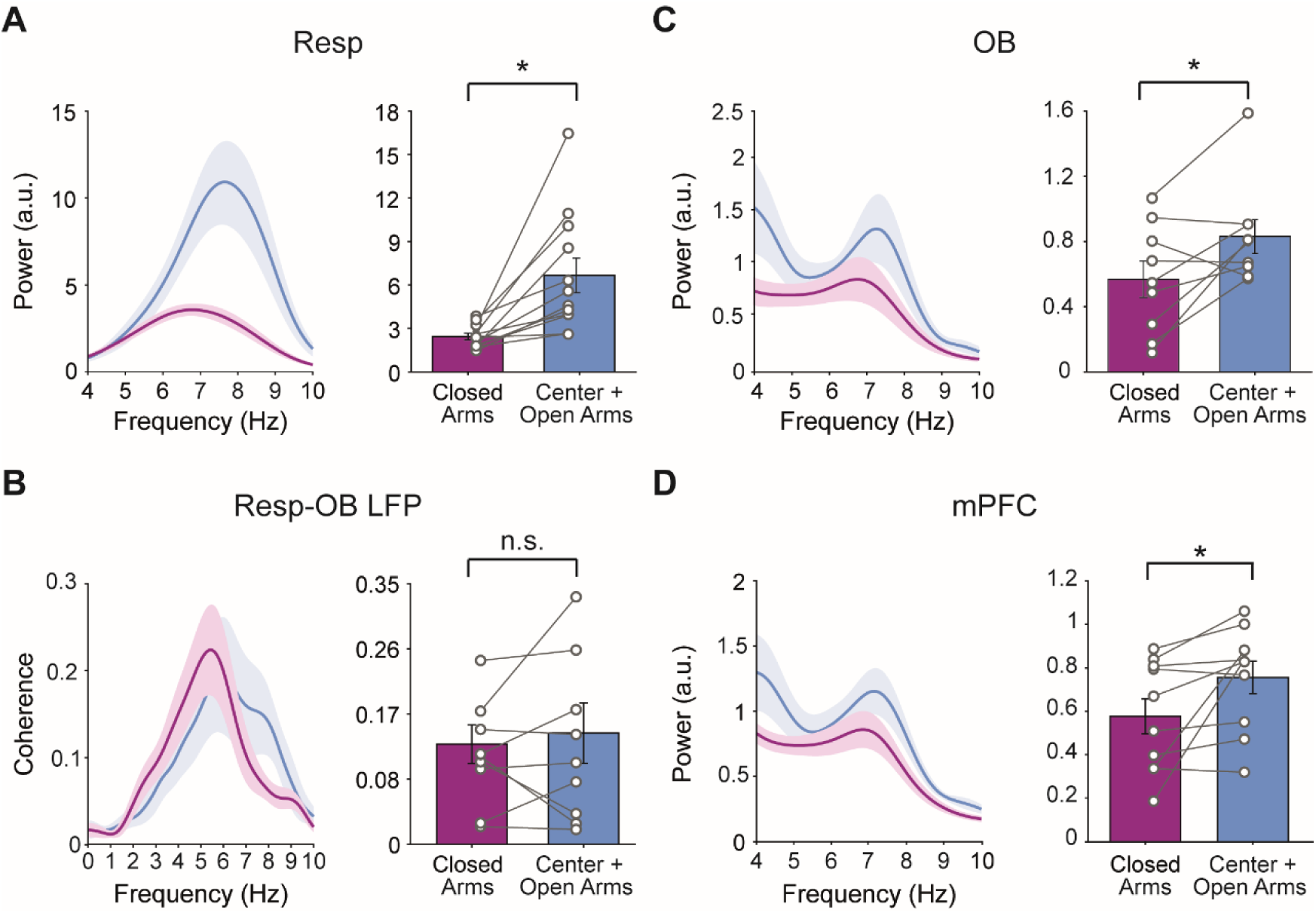
The OB and mPFC exhibit higher LFP power at fast breathing frequencies during anxious states. **(A)** Power spectrum density (left) and mean power (right) of the respiratory signal for the closed arms (pink) and center + open arms (blue). **(B)** Phase coherence spectrum between respiration and the OB LFP. **(C,D)** Power analyses as in A but for the OB (C) and MO (D) LFPs (see Figure S6 for the other mPFC regions). Notice the higher power during center + open arm exploration (*p<0.05, paired t-test) without changes in phase coherence levels (n.s., not significant). These results were obtained by considering only periods with breathing rate >4 Hz to match the breathing frequency between maze zones.

### RCOs are driven by nasal respiration

Recent studies have shown the crucial role of nasal respiration in generating RCOs, as demonstrated by their absence in tracheostomized (Yanovsky et al., 2014; Lockmann et al., 2016) and bulbectomized animals (Biskamp et al., 2017). RCOs play a significant role in facilitating long-range communication especially within slow frequency ranges (Ito et al., 2014; Lockmann and Tort, 2018; Karalis and Sirota, 2022), such as the 4 Hz rhythm seeing during freezing behavior (Moberly et al., 2018; Bagur et al., 2021). To evaluate whether both the slow and fast RCOs we observed in the anxiogenic EPM (Figures 5 and 6) are also driven by nasal respiration, we next computed bidirectional Granger causality between nasal pressure and OB or mPFC LFPs.

Our findings show that nasal respiration Granger-causes RCOs in the OB and mPFC across varying frequency ranges, exhibiting a particularly higher causal relationship at faster frequencies (Figure 7). Both individual t-tests and two-way ANOVA using directionality (LFP → Resp vs. Resp → LFP) and frequency range (0.5 – 4 Hz vs. 5 – 10 Hz) as factors revealed a statistically significant higher Granger causality for the Resp → LFP direction in all recorded regions (Figure 7 and Figure S7). Moreover, for the mPFC two-way ANOVA further revealed an effect of the frequency range (F(1,40) = 4.38, p = 0.043) as well as an interaction effect (F(1,40) = 5.71, p = 0.022), in which causality was even greater for the faster breathing in the Resp → LFP direction (Figure 7B and Figure S7). This observation is consistent with our previous results, wherein RCOs displayed heightened coherence with respiration within the faster frequency range when animals explored the center + open arms (Figure 5). In conclusion, nasal respiration drives RCOs in frontal brain regions during EPM exploration, with a particularly prominent effect for faster breathing frequencies.

**Figure 7.**
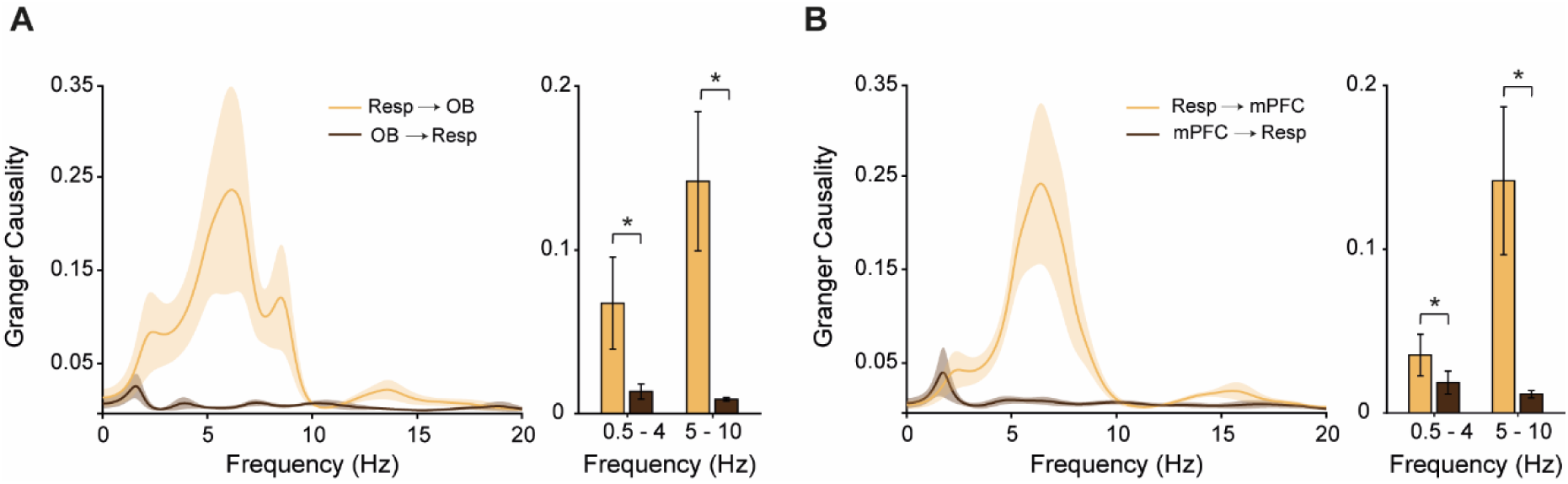
Nasal respiration drives LFP oscillations during exposition to the EPM. **(A)** Spectral Granger causality between the respiratory signal and the OB LFP during the EPM task. The right bars show the average Granger causality for each direction (Resp → LFP and LFP → Resp) separately for slow and fast respiration frequencies. **(B)** As in A but for the MO LFP. *p < 0.05, paired t-test (N = 9 – 11 sessions). See Figure S7 for the other mPFC regions.

### Breathing differentially modulates prefrontal gamma oscillations across EPM zones

It has been previously shown that the phase of the respiratory cycle modulates the amplitude of mPFC gamma oscillations (Biskamp et al., 2017; Zhong et al., 2017), suggesting an influence of breathing on the processing local information flow (González et al., 2023a, 2023b). We thus conducted comodulation analyses (Tort et al., 2010) to assess whether the different breathing regimes associated with different levels of anxiety affect the dynamics of gamma oscillations in the OB and mPFC.

We computed phase-amplitude comodulograms to assess the extent to which the power at different frequency bands of the LFP is modulated by the phase of the respiratory cycle across breathing rates. Average comodulograms were obtained for periods when the animal was in the closed arm and contrasted with those of the center + open arms (Figure 8A). Interestingly, visual inspection revealed differential modulation of prefrontal gamma oscillations by breathing depending on EPM location. During closed arm exploration, the phase of slow breathing modulated the amplitude of gamma activity ∼85 Hz, while, during center + open arm exploration, the phase of faster respiratory frequencies modulated a faster gamma activity at ∼100 Hz (Figure 8A and Figure S8).

**Figure 8.**
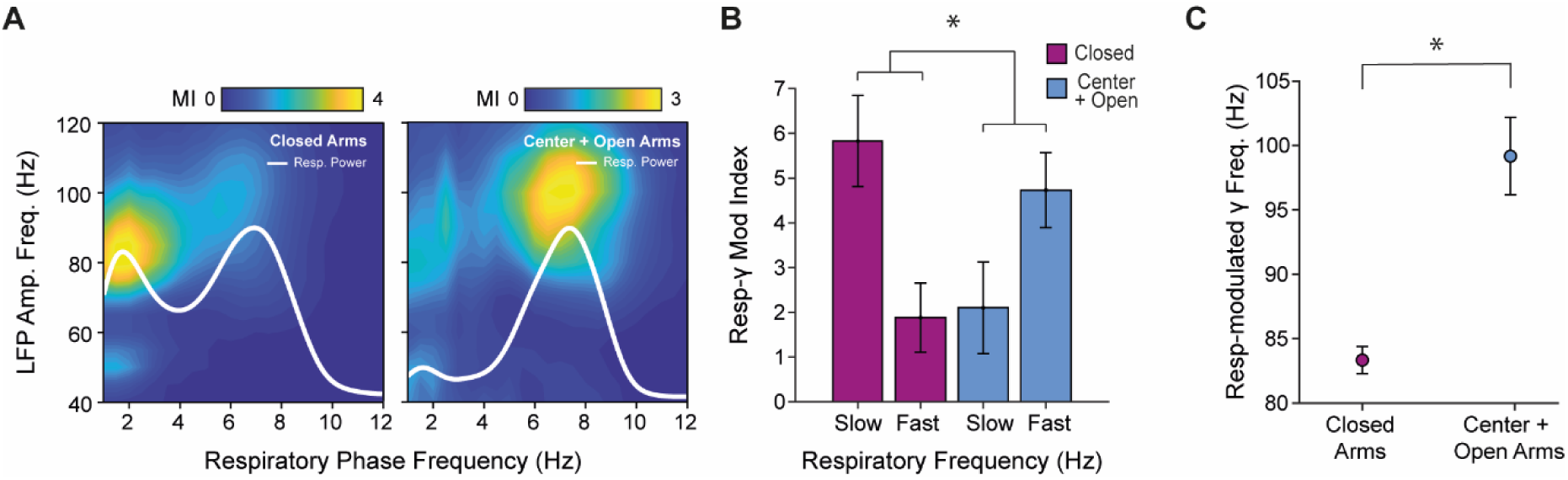
Breathing differentially modulates prefrontal gamma oscillations across EPM zones. **(A)** Average respiration – MO LFP comodulograms (n = 6 animals) for periods when the animal was in the closed arms (left) and in the center + open arms (right). Overlaid white traces show the mean respiration power spectrum. Before averaging across animals, modulation indexes were z-score normalized. See Figure S8 for the other mPFC regions. **(B)** Resp – gamma (70 – 120 Hz) normalized modulation index (n = 6 – 7 animals; *p = 0.0019, two-way ANOVA interaction between maze region and respiratory frequency). **(C)** Modulated gamma frequency by breathing in the closed and center + open arms by slow (0.5 – 4 Hz) and fast (5 – 10 Hz) respiratory frequencies, respectively. *p = 0.0021, paired t-test.

We subsequently performed statistical analysis to confirm these observations. For each session, we obtained from the comodulogram the maximal modulation index within the gamma frequency (70 – 120 Hz) for the two breathing regimes (0.5 – 4 Hz and 5–10 Hz). Two-way ANOVA revealed a significant interaction between maze region and respiratory frequency (F(1,20) = 12.77, p = 0.0019), in which gamma oscillations were more modulated by the slower respiratory frequency when the animal was in the closed arms, whereas the opposite effect was observed during periods in the center + open arms (Figure 8B). Additionally, after obtaining the gamma frequency of peak phase-amplitude modulation from the comodulogram, we found a statistically significantly faster gamma activity during center + open arm compared to closed arm exposition (t(5) = 5.84, p = 0.0021, t-test; Figure 8C). Therefore, our results demonstrate a differential modulation of prefrontal gamma oscillations by breathing depending on anxious state.

## Discussion

In our study, we have shown that rats explore more the center + open arms and perform more risk assessment behavior during the initial minutes of the EPM task (Figure 2). EPM exploration was accompanied by distinct respiratory patterns: a slower breathing frequency (∼ 0.5 – 4 Hz) predominated during periods spent in the closed arms, while a faster frequency (∼ 5 – 10 Hz) was observed during exploration of the center + open arms (Figure 3). Importantly, the increase in breathing rate during center + open arms exploration was specifically related to the maze zone rather than to changes in animal speed (Figure 4), so that anxiety-like states led to higher respiratory rates independently of locomotor activity. Upon closer examination of the electrophysiological recordings, we identified RCOs at different frequency bands during the task (Figure 5), mirroring the bimodal distribution observed in respiratory frequency. Moreover, RCO power was notably higher during the center + open arm exploration associated with fast breathing (> 4 Hz); RCOs also exhibited greater coherence with respiration at faster frequencies during these periods (Figures 5 and 6). Additionally, we observed that it is the nasal respiration that is driving the RCOs in frontal brain regions during EPM exploration with a greater effect at faster breathing frequencies (Figure 7). Finally, comodulation analyses revealed that the amplitude of prefrontal gamma oscillations is modulated by the breathing cycle, and the frequency of the modulated gamma activity is dependent on anxiety levels (Figure 8).

The elevated plus maze is widely employed for assessing anxiety-like behavior in rodents due to its cost-effectiveness, efficiency, and ethological relevance (Carobrez and Bertoglio, 2005). The EPM was initially designed to investigate anxiolytic drugs, in which increased or decreased exploration of the open arms respectively indicates anxiolytic or anxiogenic effects (Lister, 1987). While traditionally used for pharmacological purposes, our study aimed to investigate different anxiety levels displayed within an EPM session. Since previous findings have shown increased anxiety during open arms exploration (Pellow et al., 1985; Lister, 1987; Okonogi et al., 2018), we divided our analyses according to the EPM zone: the closed arms, representing relatively protected areas associated with less anxiety-like behavior, and the center + open arms, which evoke heightened anxiety levels.

It is important to note that behavior in the EPM is commonly assessed with only one session because repeated exposures result in gradual habituation to the maze and a reduction in the anxiogenic effect of the open arms (File, 1993; Holmes and Rodgers, 1998); moreover, EPM sessions usually last 5 minutes (Walf and Frye, 2007). In our work, to maximize data yield within the maze and potentially increase the time spent exploring the open arms, we exposed the animals to two EPM sessions of 10 minutes. Interestingly, in the current dataset the behavior of the animals did not differ between sessions (Figures 2, S2 and S4), which allowed for a combined analysis of the electrophysiological and breathing recordings along with behavior. In both sessions, we observed that the animals exhibited SAP, higher locomotion and explored more the center + open arms in the initial minutes (Figure 2A,C,D), consistent with a faster respiratory rate at the beginning of the task (Figure 3C). Moreover, similar to breathing rate, these behaviors decreased along the task while the time spent in the closed arms increased, as inferred from the minute-by-minute analyses.

In rodents, breathing can occur within different frequency ranges; it is usually slower during resting states (∼0.5–4 Hz) and faster during active exploration and sniffing (∼7–14 Hz) (Courtiol et al., 2011; Wachowiak, 2011; Rojas-Líbano et al., 2014; Sirotin et al., 2014). By measuring EMG, heart and respiratory rates, Okonogi et al. (2018) have previously shown that mice exhibit both low and high peripheral activity in the closed arms, while only exhibiting high activity in the open arms. Their findings match ours in which both slow and fast respiratory frequencies could be observed in the closed arms while fast breathing predominated in the open arms.

Since variations in breathing rate also relate to animal speed (Bramble and Carrier, 1983; Nguyen Chi et al., 2016), here we analyzed locomotion in the maze to disentangle its influence from that of putative anxiety states. Our results showed that, during closed arm exploration, locomotion speed was indeed a main factor influencing respiratory rate, with slower breathing at ∼3 Hz observed at 2.5 cm/s average speed, while a faster breathing rate of ∼ 5 Hz occurred for the average speed of 18 cm/s. Interestingly, however, during center + open arm exploration, we observed high respiratory rates of 6 Hz for the lowest analyzed speed bin, which mildly increased to 6.5 Hz for the fastest speed. Therefore, although there were times in the EPM when locomotion could influence breathing, in the center + open arms animals exhibited increased respiratory frequency regardless of their speed (Figure 4). Our results are consistent with previous studies showing that animals exposed to aversive or anxiogenic stimuli increase the respiratory rate (Carnevali et al., 2013; Okonogi et al., 2018), supporting the idea that breathing varies depending on behavioral state or emotional valence (Ashhad et al., 2022). We conclude that the anxiety-like state associated with center + open arm exploration is a main factor influencing respiratory rate in the EPM.

By analyzing synchronized behavioral, respiratory and LFP activities, our study revealed the presence of field potential oscillations entrained by respiration (RCOs) in the OB and mPFC during the EPM task, which exhibited a prominent bimodal power-frequency profile, mirroring the distribution of respiratory rates. The appearance of RCOs in slow (∼ 0.5 – 4 Hz) and fast (∼ 5 – 10 Hz) frequencies during the periods in the closed arms is consistent with the findings of Okonogi et al. (2018), in which the animals displayed not only low and high peripheral activity but also different levels of delta and theta power in the closed arms. Interestingly, we found that RCOs have higher power and heightened coherence with breathing in the theta frequency range (5 – 10 Hz) when the animals were in the center + open arms (Figure 5 and 6). This result is again consistent with Okonogi et al. (2018) who observed greater theta power while mice explored the open arm.

While anxiety-like behaviors have been associated with the appearance of theta 2 oscillations (Adhikari et al., 2010; Mikulovic et al., 2018; Totty and Maren, 2022; Mooziri et al., 2024), we have shown that theta rhythms can be confounded with RCOs during periods of faster breathing (Tort et al., 2018a, 2018b). To mitigate the appearance of hippocampal and parahippocampal theta (Brankačk et al., 1993; Sirota et al., 2008), in the present work we placed our reference electrode over the parietal cortex to subtract its prominent theta rhythms from our recording electrodes. With this approach, we unveiled RCOs in anxiety-like states with stronger power and coherence to breathing in the same frequency range as theta during open arm exploration; moreover, Granger causality analysis showed that the RCOs are strongly driven by nasal respiration at this frequency range. Taken together, our results suggest that previous findings reporting theta oscillations during anxiety-like behaviors may have actually observed RCO activity, which was not identified as such due to the absence of concomitant respiration recordings. This may have been the case in the recent study by Mooziri et al. (2024), who reported synchronization between the OB and mPFC at theta frequency during EPM exploration – a finding consistent with our present results. In any event, our study underscores the importance of considering respiratory dynamics when interpreting neural oscillatory patterns associated with emotional states.

Previous studies have proposed that RCOs emerge in the frontal brain during a myriad of emotional states, including positive and negative ones (Folschweiller and Sauer, 2021, 2023). In particular, RCOs in the delta frequency range (2 – 4 Hz) have been closely associated with the initiation and maintenance of freezing behavior (Karalis et al., 2016; Moberly et al., 2018; Bagur et al., 2021). Our results now show that frontal RCOs also appear at faster respiratory rates during anxiety-like states, including the theta range.

Gamma oscillations in the range of 30 – 160 Hz are believed to play a pivotal role in different cognitive processes (Buzsáki and Wang, 2012). The functional significance of gamma oscillations extends beyond their isolated occurrence and may relate to their coupling with slower rhythmic activities, such as theta (Sirota et al., 2008; Tort et al., 2009; Belluscio et al., 2012; Lisman and Jensen, 2013) or respiration (Zhong et al., 2017; Cavelli et al., 2020; Sheriff et al., 2021; González et al., 2023b). Several reports have shown that the phase of RCOs modulates the amplitude of gamma oscillations in the mPFC (Biskamp et al., 2017; Tort et al., 2018b; Karalis and Sirota, 2022; Basha et al., 2023; Folschweiller and Sauer, 2023), suggesting their potential involvement in processing local multimodal information flow in these regions across different species (González et al., 2023a).

In order to gain further insights into this phenomenon, here we measured phase-amplitude coupling between prefrontal gamma activity and breathing during the EPM task. Our results showed a differential modulation of gamma by breathing depending on emotional/behavioral state. Namely, during periods in the closed arms (putative low anxiety state), slow breathing modulates a gamma frequency around 85 Hz, while during periods in the center + open arms (putative high anxiety states), faster breathing modulates a faster gamma around 100 Hz. Our results thus show that not only the breathing frequency but also the frequency of respiration-modulated prefrontal gamma oscillations increase during anxiety. Moreover, they are consistent with recent studies showing that, in addition to phase-amplitude entrainment, respiration and brain activity also couple through frequency-frequency relations (Hammer et al., 2021; Tort et al., 2021). While the functional significance of these findings remains to be unveiled, it may be that faster rhythmic activity would allow faster computations and decision-making to be performed, as required to cope with perceived threats.

Brain regions commonly implicated in fearful experiences include the mPFC, hippocampus and amygdala (Calhoon and Tye, 2015; Felix-Ortiz et al., 2016; Mikulovic et al., 2018; Jacobs and Moghaddam, 2021; Shi et al., 2023). These regions activate and synchronize their activity during freezing and anxiety-like behaviors (Adhikari et al., 2010, 2011; Jacinto et al., 2016). Interestingly, anxiolytic drugs diminish both anxious behavior and theta frequency in the mPFC and hippocampus (Zhu and McNaughton, 1994; McNaughton et al., 2007; Yeung et al., 2012). In addition to these regions, the present findings, along with other studies (Moberly et al., 2018; Bagur et al., 2021; Karalis and Sirota, 2022), suggest the involvement of the OB in coordinating pathways associated with fear and anxiety through mediating RCOs to downstream regions (see also Mooziri et al., 2024). This idea is supported by the existence of direct projections from the olfactory system to the prelimbic prefrontal cortex (Clugnet and Price, 1987), along with the fact that disruption of olfactory inputs leads to depressive behaviors (Song and Leonard, 2005) and affect freezing expression (Bagur et al., 2021). In all, our work provides evidence for a pronounced influence of breathing on the activity of prefrontal cortex networks during anxious states. Moreover, it underscores the role of breathing as a model for a general mechanism through which the body influences brain functioning (Heck et al., 2017, 2022).

## Acknowledgments

This work was supported by the Brazilian National Council for Scientific and Technological Development (CNPq), the Brazilian Coordination for the Improvement of Higher Education Personnel (CAPES), and the Alexander von Humboldt Foundation.

## Supplementary Material

**Supplementary Figure S1.**
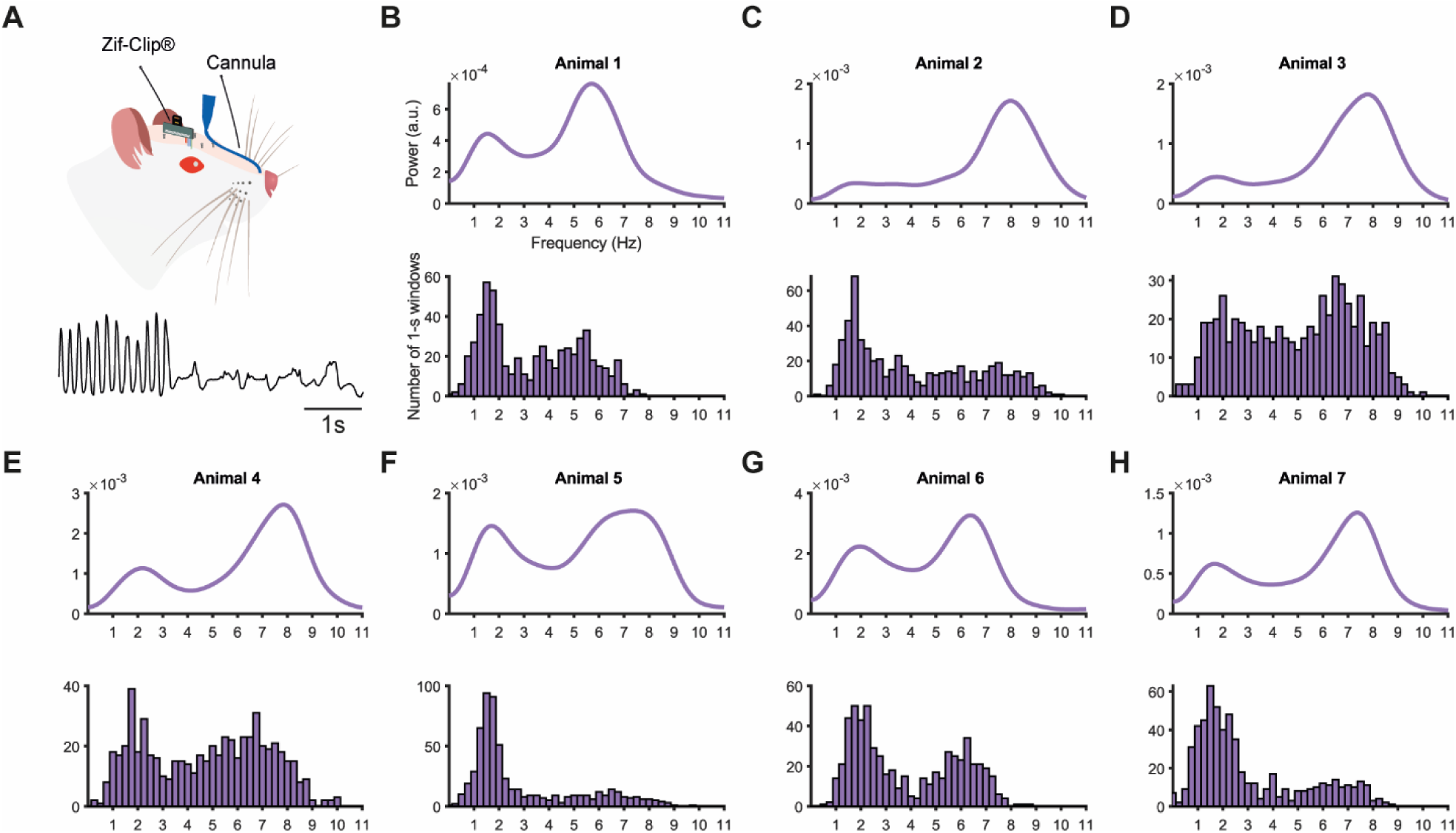
Bimodal distribution of respiratory frequency in the EPM. **(A)** Illustration of an animal implanted with a pressure cannula (top) and a respiratory trace example (bottom). **(B-H)** Power spectrum (top) and respiratory rate histograms (bottom) for each animal. Notice a bimodal distribution, with animals tending to breath at either slower (∼ 0.5–4 Hz) or faster (∼ 5–10 Hz) respiratory rates. Note that a higher power for faster frequencies reflects a higher air pressure amplitude and does not necessarily indicate a higher occurrence of fast respiration frequencies.

**Supplementary Figure S2.**
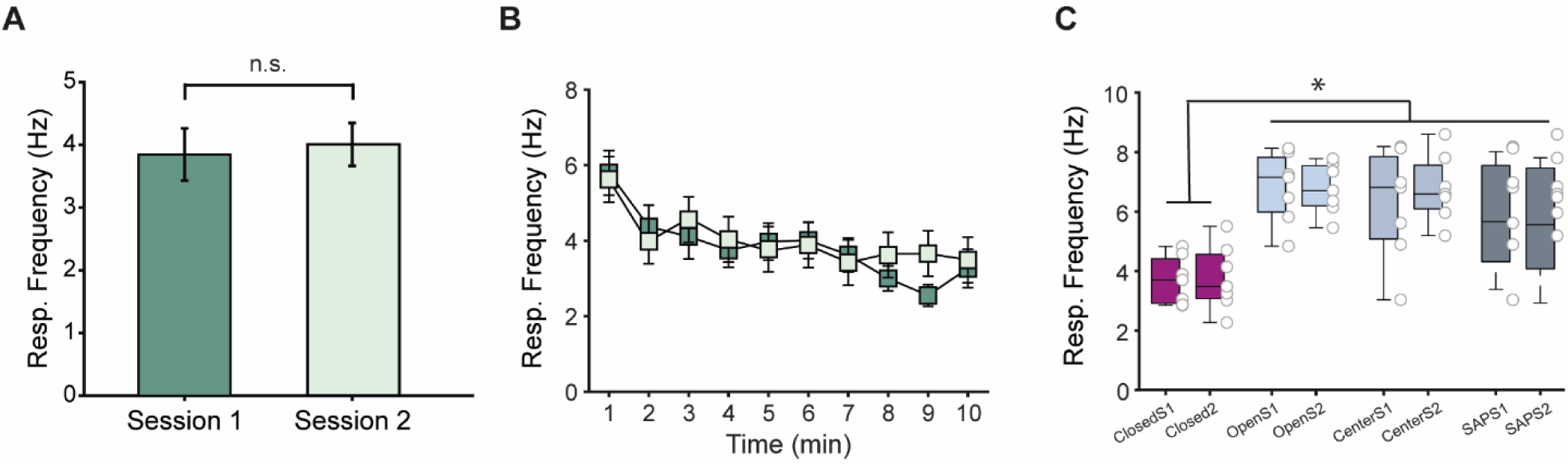
Similar respiratory frequencies across sessions in the EPM. **(A,B)** Comparison of the mean respiratory frequency (A) and their minute-by-minute time course in the EPM (B) between sessions 1 and 2. **(C)** Respiratory frequency in the different zones of the maze and during SAP events (N = 7 animals) in the two experimental sessions. *p<0.0001, repeated measures ANOVA followed by the Tukey-Kramer test.

**Supplementary Figure S3.**
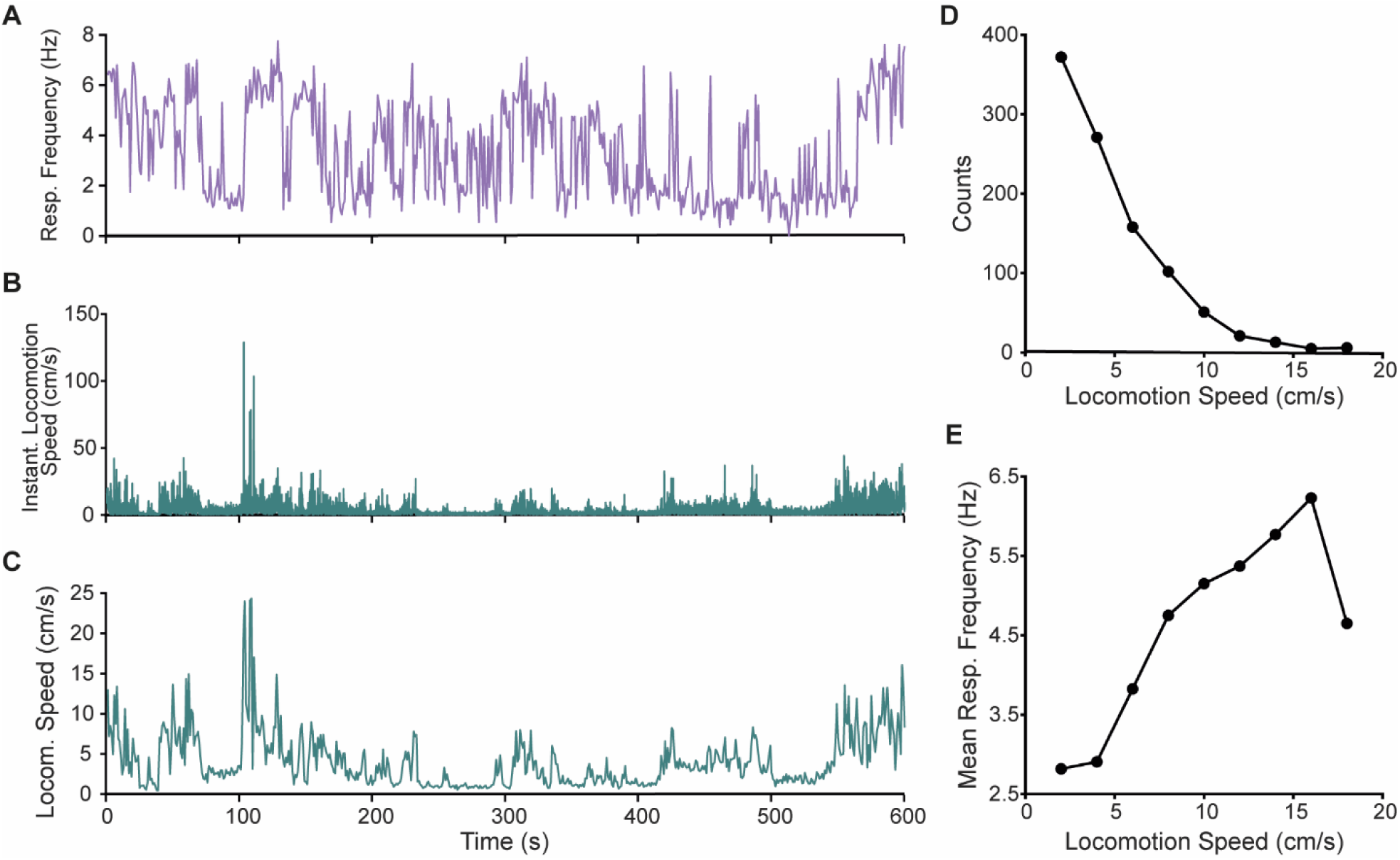
Respiratory frequency is related to locomotion speed in the EPM. **(A,B)** Respiratory frequency in 1-second windows (A) and instantaneous locomotion speed estimated at each video frame (B). **(C)** Average speed in non-overlapping 1-second windows (i.e., as used to compute the respiratory frequency, so that the two times series can be matched). **(D)** Histogram counts of the average locomotion speed shown in C. **(E)** Mean respiratory frequency as a function of the locomotion speed for the data in A and C. Results in A-E are from a representative animal. Speed was estimated from the *x* and *y* coordinates of the animal’s head.

**Supplementary Figure S4.**
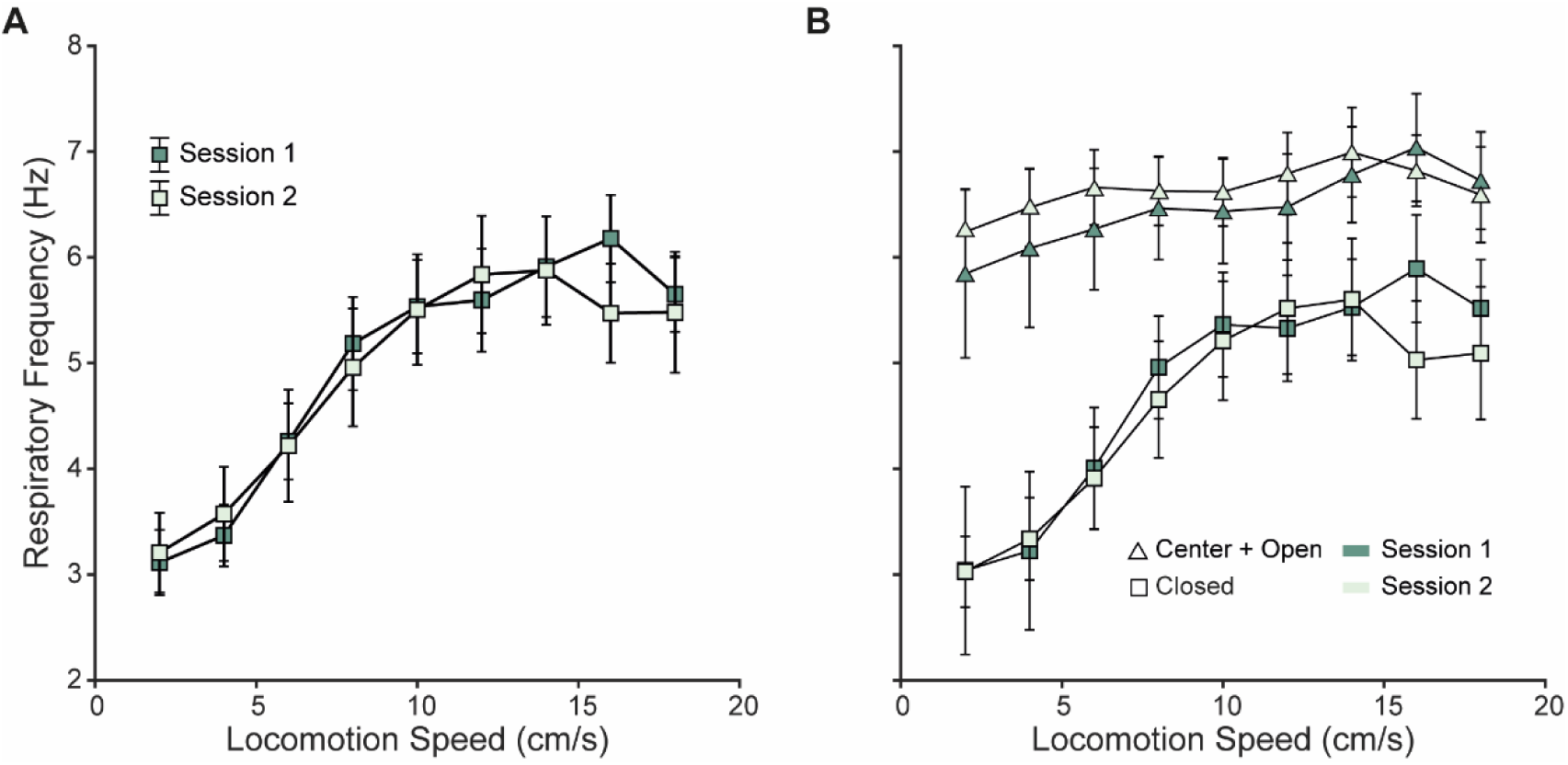
Similar relationship between respiratory frequency and locomotion speed across sessions in the EPM. **(A)** Average respiratory frequency (± SEM) at different locomotion speeds in the two EPM sessions (N = 7 animals). **(B)** Average respiratory frequency as a function of speed plotted separately for closed arms and center & open arms for each EPM session. Speed was estimated from the head *x* and *y* coordinates.

**Supplementary Figure S5.**
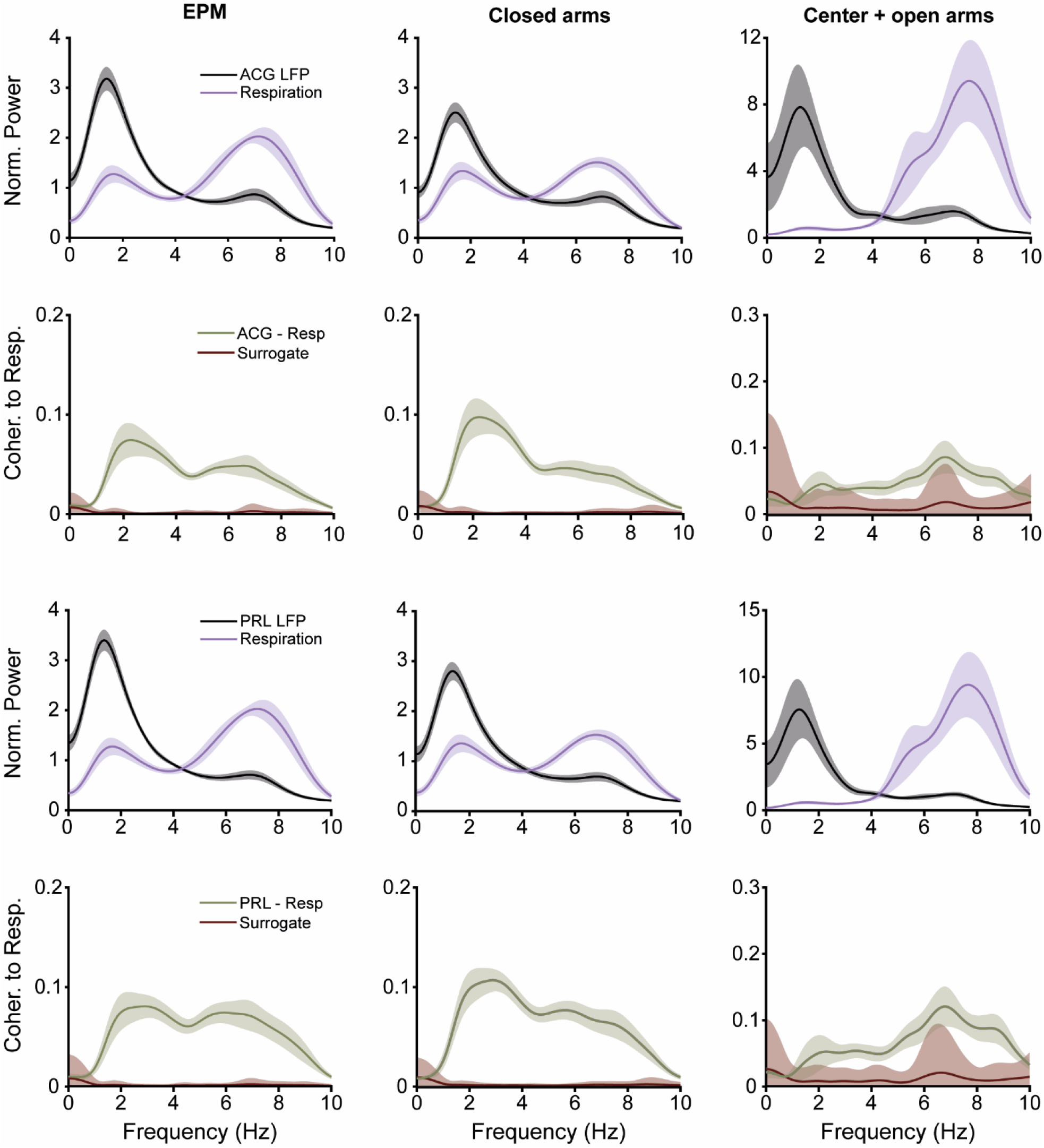
Respiration-coupled LFP oscillations in the EPM. Panels show the same as in Figure 5 for two additional regions (ACG and PRL).

**Supplementary Figure S6.**
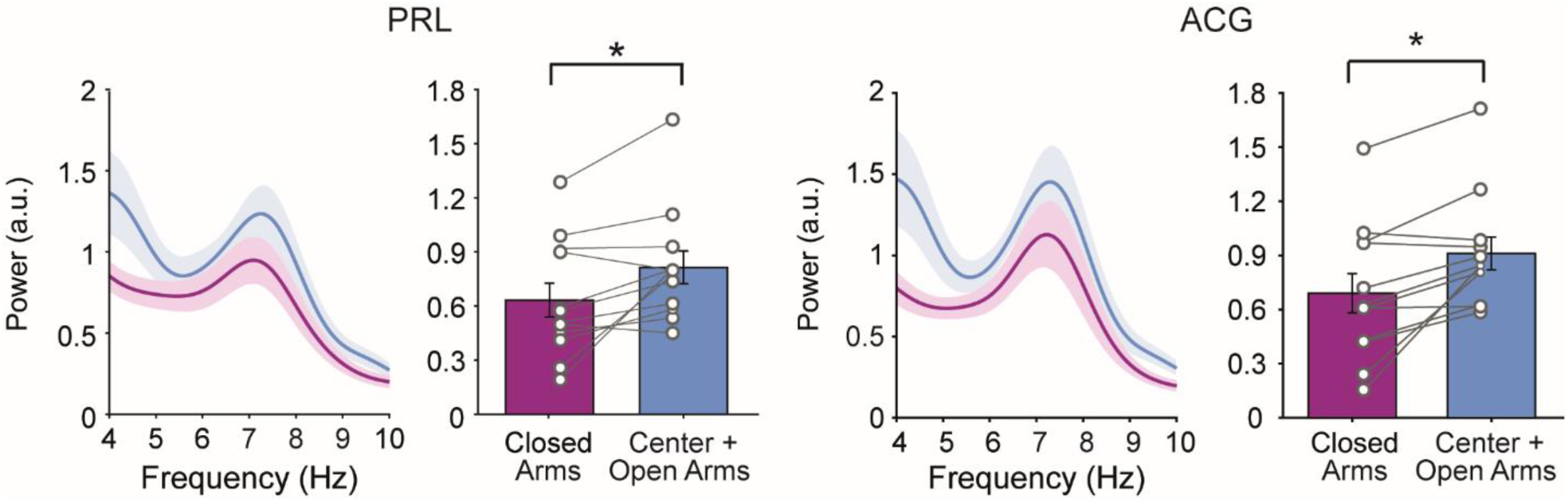
The mPFC exhibits higher LFP power at fast breathing frequencies during anxious states. Panels show the same as in Figure 6 but for two additional regions (PRL and ACG). *p< 0.05, paired t-test.

**Supplementary Figure S7.**
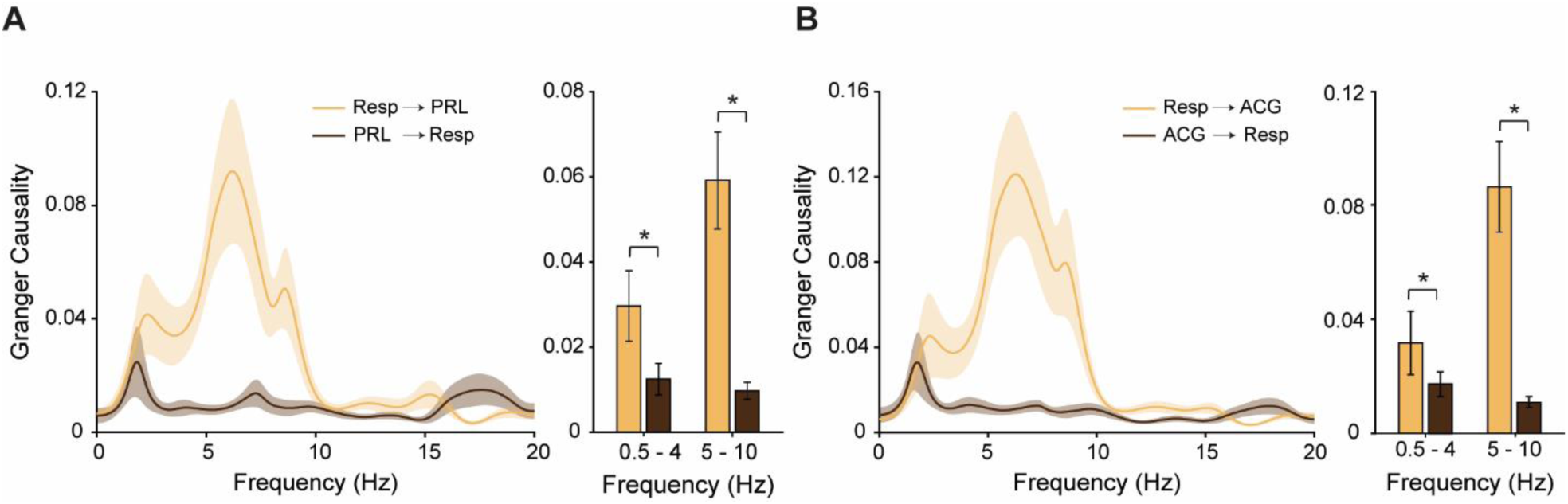
Respiration drives LFP oscillations during exposition to the EPM. Panels show the same as in Figure 7 but for two additional regions (ACG and PRL). *p< 0.05, paired t-test (N = 13 sessions).

**Supplementary Figure S8.**
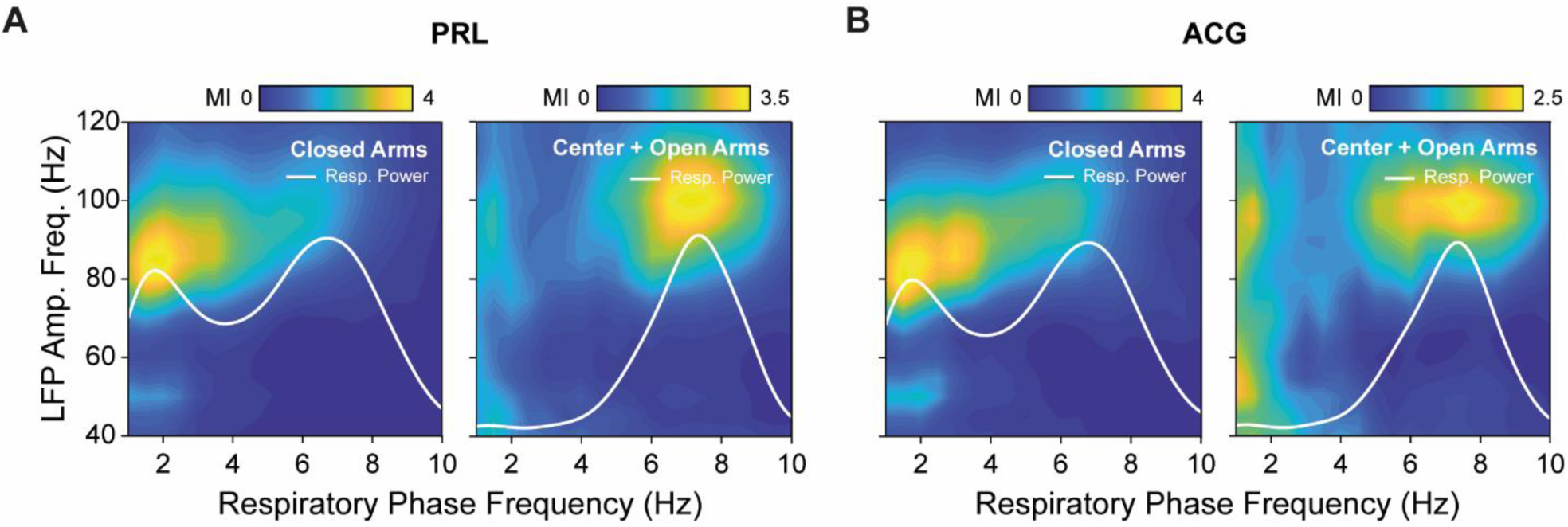
Respiration modulates gamma oscillations in the mPFC and OB. **(A)** Mean respiration - LFP comodulograms for periods when the animal was exposed to the closed arms (left) and to the center + open arms (right) in the PRL. **(B)** Same as in A but for the ACG. White traces show respiration power.

## Notes

### Competing Interest Statement

The authors have declared no competing interest.

## References

Adhikari A, Topiwala MA, Gordon JA (2010) Synchronized Activity between the Ventral Hippocampus and the Medial Prefrontal Cortex during Anxiety. Neuron 65:257–269.

Adhikari A, Topiwala MA, Gordon JA (2011) Single Units in the Medial Prefrontal Cortex with Anxiety-Related Firing Patterns Are Preferentially Influenced by Ventral Hippocampal Activity. Neuron 71:898–910.

Allen M, Varga S, Heck DH (2023) Respiratory rhythms of the predictive mind. Psychol Rev 130:1066–1080.

Ashhad S, Kam K, Del Negro CA, Feldman JL (2022) Breathing Rhythm and Pattern and Their Influence on Emotion. Annu Rev Neurosci 45:223–247.

Bagur S, Lefort JM, Lacroix MM, de Lavilléon G, Herry C, Chouvaeff M, Billand C, Geoffroy H, Benchenane K (2021) Breathing-driven prefrontal oscillations regulate maintenance of conditioned-fear evoked freezing independently of initiation. Nat Commun 12:2605.

Barnett L, Seth AK (2014) The MVGC multivariate Granger causality toolbox: a new approach to Granger-causal inference. J Neurosci Methods 223:50–68.

Basha D, Chauvette S, Sheroziya M, Timofeev I (2023) Respiration organizes gamma synchrony in the prefronto-thalamic network. Sci Rep 13:8529.

Belluscio MA, Mizuseki K, Schmidt R, Kempter R, Buzsáki G (2012) Cross-frequency phase-phase coupling between θ and γ oscillations in the hippocampus. J Neurosci 32:423–435.

Biskamp J, Bartos M, Sauer J-F (2017) Organization of prefrontal network activity by respiration-related oscillations. Sci Rep 7:45508.

Brændholt M, Kluger DS, Varga S, Heck DH, Gross J, Allen MG (2023) Breathing in waves: Understanding respiratory-brain coupling as a gradient of predictive oscillations. Neurosci Biobehav Rev 152:105262.

Bramble DM, Carrier DR (1983) Running and breathing in mammals. Science 219:251–256.

Brankačk J, Stewart M, Fox SE (1993) Current source density analysis of the hippocampal theta rhythm: associated sustained potentials and candidate synaptic generators. Brain Res 615:310–327.

Buzsáki G, Draguhn A (2004) Neuronal oscillations in cortical networks. Science 304:1926–1929.

Buzsáki G, Wang X-J (2012) Mechanisms of Gamma Oscillations. Annu Rev Neurosci 35:203–225.

Calhoon GG, Tye KM (2015) Resolving the neural circuits of anxiety. Nat Neurosci 18:1394–1404.

Carnevali L, Sgoifo A, Trombini M, Landgraf R, Neumann ID, Nalivaiko E (2013) Different patterns of respiration in rat lines selectively bred for high or low anxiety. PloS One 8:e64519.

Carobrez AP, Bertoglio LJ (2005) Ethological and temporal analyses of anxiety-like behavior: The elevated plus-maze model 20 years on. Neurosci Biobehav Rev 29:1193–1205.

Cavelli M, Castro-Zaballa S, Gonzalez J, Rojas-Líbano D, Rubido N, Velásquez N, Torterolo P (2020) Nasal respiration entrains neocortical long-range gamma coherence during wakefulness. Eur J Neurosci 51:1463–1477.

Cenier T, David F, Litaudon P, Garcia S, Amat C, Buonviso N (2009) Respiration-gated formation of gamma and beta neural assemblies in the mammalian olfactory bulb. Eur J Neurosci 29:921– 930.

Clugnet M-C, Price JL (1987) Olfactory input to the prefrontal cortex in the rat. Ann N Y Acad Sci 510:231–235.

Courtiol E, Hegoburu C, Litaudon P, Garcia S, Fourcaud-Trocmé N, Buonviso N (2011) Individual and synergistic effects of sniffing frequency and flow rate on olfactory bulb activity. J Neurophysiol 106:2813–2824.

Delorme A, Makeig S (2004) EEGLAB: an open source toolbox for analysis of single-trial EEG dynamics including independent component analysis. J Neurosci Methods 134:9–21.

Felix-Ortiz AC, Burgos-Robles A, Bhagat ND, Leppla CA, Tye KM (2016) Bidirectional modulation of anxiety-related and social behaviors by amygdala projections to the medial prefrontal cortex. Neuroscience 321:197–209.

File SE (1993) The interplay of learning and anxiety in the elevated plus-maze. Behav Brain Res 58:199–202.

Folschweiller S, Sauer J-F (2021) Respiration-Driven Brain Oscillations in Emotional Cognition. Front Neural Circuits 15:761812.

Folschweiller S, Sauer J-F (2023) Behavioral State-Dependent Modulation of Prefrontal Cortex Activity by Respiration. J Neurosci 43:4795–4807.

Girin B, Juventin M, Garcia S, Lefèvre L, Amat C, Fourcaud-Trocmé N, Buonviso N (2021) The deep and slow breathing characterizing rest favors brain respiratory-drive. Sci Rep 11:7044.

González J, Cavelli M, Mondino A, Castro-Zaballa S, Brankačk J, Draguhn A, Torterolo P, Tort ABL (2023a) Breathing modulates gamma synchronization across species. Pflugers Arch 475:49– 63.

González J, Torterolo P, Tort ABL (2023b) Mechanisms and functions of respiration-driven gamma oscillations in the primary olfactory cortex. eLife 12:e83044.

Goutagny R, Jackson J, Williams S (2009) Self-generated theta oscillations in the hippocampus. Nat Neurosci 12:1491–1493.

Grosmaitre X, Santarelli LC, Tan J, Luo M, Ma M (2007) Dual functions of mammalian olfactory sensory neurons as odor detectors and mechanical sensors. Nat Neurosci 10:348–354.

Hammer M, Schwale C, Brankačk J, Draguhn A, Tort ABL (2021) Theta-gamma coupling during REM sleep depends on breathing rate. Sleep 44:zsab189.

Heck DH, Correia BL, Fox MB, Liu Y, Allen M, Varga S (2022) Recent insights into respiratory modulation of brain activity offer new perspectives on cognition and emotion. Biol Psychol 170:108316.

Heck DH, McAfee SS, Liu Y, Babajani-Feremi A, Rezaie R, Freeman WJ, Wheless JW, Papanicolaou AC, Ruszinkó M, Sokolov Y, Kozma R (2017) Breathing as a fundamental rhythm of brain function. Front Neural Circuits 10:115.

Hofmann SG, Gómez AF (2017) Mindfulness-Based Interventions for Anxiety and Depression. Psychiatr Clin North Am 40:739–749.

Holmes A, Rodgers RJ (1998) Responses of Swiss–Webster Mice to Repeated Plus-Maze Experience Further Evidence for a Qualitative Shift in Emotional State? Pharmacol Biochem Behav 60:473–488.

Ito J, Roy S, Liu Y, Cao Y, Fletcher M, Lu L, Boughter JD, Grün S, Heck DH (2014) Whisker barrel cortex delta oscillations and gamma power in the awake mouse are linked to respiration. Nat Commun 5:3572.

Jacinto LR, Cerqueira JJ, Sousa N (2016) Patterns of Theta Activity in Limbic Anxiety Circuit Preceding Exploratory Behavior in Approach-Avoidance Conflict. Front Behav Neurosci 10:171.

Jacobs DS, Moghaddam B (2021) Medial prefrontal cortex encoding of stress and anxiety. Int Rev Neurobiol 158:29–55.

Karalis N, Dejean C, Chaudun F, Khoder S, Rozeske RR, Wurtz H, Bagur S, Benchenane K, Sirota A, Courtin J, Herry C (2016) 4-Hz oscillations synchronize prefrontal-amygdala circuits during fear behavior. Nat Neurosci 19:605–612.

Karalis N, Sirota A (2022) Breathing coordinates cortico-hippocampal dynamics in mice during offline states. Nat Commun 13:467.

Lesting J, Narayanan RT, Kluge C, Sangha S, Seidenbecher T, Pape H-C (2011) Patterns of coupled theta activity in amygdala-hippocampal-prefrontal cortical circuits during fear extinction. PloS One 6:e21714.

Lisman JE, Jensen O (2013) The Theta-Gamma Neural Code. Neuron 77:1002–1016.

Lister RG (1987) The use of a plus-maze to measure anxiety in the mouse. Psychopharmacology (Berl) 92:180–185.

Lockmann ALV, Laplagne DA, Leão RN, Tort ABL (2016) A respiration-coupled rhythm in the rat hippocampus independent of theta and slow oscillations. J Neurosci 36:5338–5352.

Lockmann ALV, Tort ABL (2018) Nasal respiration entrains delta-frequency oscillations in the prefrontal cortex and hippocampus of rodents. Brain Struct Funct 223:1–3.

Masaoka Y, Homma I (1997) Anxiety and respiratory patterns: their relationship during mental stress and physical load. Int J Psychophysiol 27:153–159.

McNaughton N, Kocsis B, Hajós M (2007) Elicited hippocampal theta rhythm: a screen for anxiolytic and procognitive drugs through changes in hippocampal function? Behav Pharmacol 18:329– 346.

Mikulovic S, Restrepo CE, Siwani S, Bauer P, Pupe S, Tort ABL, Kullander K, Leão RN (2018) Ventral hippocampal OLM cells control type 2 theta oscillations and response to predator odor. Nat Commun 9:3638.

Moberly AH, Schreck M, Bhattarai JP, Zweifel LS, Luo W, Ma M (2018) Olfactory inputs modulate respiration-related rhythmic activity in the prefrontal cortex and freezing behavior. Nat Commun 9:1528.

Mooziri M, Samii Moghaddam A, Mirshekar MA, Raoufy MR (2024) Olfactory bulb-medial prefrontal cortex theta synchronization is associated with anxiety. Sci Rep 14:12101.

Nagayama S, Homma R, Imamura F (2014) Neuronal organization of olfactory bulb circuits. Front Neural Circuits 8:98.

Nath T, Mathis A, Chen AC, Patel A, Bethge M, Mathis MW (2019) Using DeepLabCut for 3D markerless pose estimation across species and behaviors. Nat Protoc 14:2152–2176.

Nguyen Chi V, Müller C, Wolfenstetter T, Yanovsky Y, Draguhn A, Tort ABL, Brankačk J (2016) Hippocampal respiration-driven rhythm distinct from theta oscillations in awake mice. J Neurosci 36:162–177.

Okonogi T, Nakayama R, Sasaki T, Ikegaya Y (2018) Characterization of Peripheral Activity States and Cortical Local Field Potentials of Mice in an Elevated Plus Maze Test. Front Behav Neurosci 12:62.

Pellow S, Chopin P, File SE, Briley M (1985) Validation of open : closed arm entries in an elevated plus-maze as a measure of anxiety in the rat. J Neurosci Methods 14:149–167.

Rojas-Líbano D, Frederick DE, Egaña JI, Kay LM (2014) The olfactory bulb theta rhythm follows all frequencies of diaphragmatic respiration in the freely behaving rat. Front Behav Neurosci 8.

Sheriff A, Pandolfi G, Nguyen VS, Kay LM (2021) Long-Range Respiratory and Theta Oscillation Networks Depend on Spatial Sensory Context. J Neurosci 41:9957–9970.

Shi H-J, Wang S, Wang X-P, Zhang R-X, Zhu L-J (2023) Hippocampus: Molecular, Cellular, and Circuit Features in Anxiety. Neurosci Bull 39:1009–1026.

Sirota A, Montgomery S, Fujisawa S, Isomura Y, Zugaro M, Buzsáki G (2008) Entrainment of neocortical neurons and gamma oscillations by the hippocampal theta rhythm. Neuron 60:683–697.

Sirotin YB, Costa ME, Laplagne DA (2014) Rodent ultrasonic vocalizations are bound to active sniffing behavior. Front Behav Neurosci 8 Available at: http://www.ncbi.nlm.nih.gov/pmc/articles/PMC4235378/ [Accessed August 24, 2015].

Song C, Leonard BE (2005) The olfactory bulbectomised rat as a model of depression. Neurosci Biobehav Rev 29:627–647.

Stujenske JM, Likhtik E, Topiwala MA, Gordon JA (2014) Fear and safety engage competing patterns of theta-gamma coupling in the basolateral amygdala. Neuron 83:919–933.

Tort ABL, Brankačk J, Draguhn A (2018a) Respiration-Entrained Brain Rhythms Are Global but Often Overlooked. Trends Neurosci 41:186–197.

Tort ABL, Hammer M, Zhang J, Brankačk J, Draguhn A (2021) Temporal Relations between Cortical Network Oscillations and Breathing Frequency during REM Sleep. J Neurosci 41:5229–5242.

Tort ABL, Komorowski R, Eichenbaum H, Kopell N (2010) Measuring phase-amplitude coupling between neuronal oscillations of different frequencies. J Neurophysiol 104:1195–1210.

Tort ABL, Komorowski RW, Manns JR, Kopell NJ, Eichenbaum H (2009) Theta-gamma coupling increases during the learning of item-context associations. Proc Natl Acad Sci U S A 106:20942–20947.

Tort ABL, Ponsel S, Jessberger J, Yanovsky Y, Brankačk J, Draguhn A (2018b) Parallel detection of theta and respiration-coupled oscillations throughout the mouse brain. Sci Rep 8:6432.

Totty MS, Maren S (2022) Neural Oscillations in Aversively Motivated Behavior. Front Behav Neurosci 16:936036.

Wachowiak M (2011) All in a sniff: olfaction as a model for active sensing. Neuron 71:962–973.

Walf AA, Frye CA (2007) The use of the elevated plus maze as an assay of anxiety-related behavior in rodents. Nat Protoc 2:322–328.

Yanovsky Y, Ciatipis M, Draguhn A, Tort ABL, Brankačk J (2014) Slow oscillations in the mouse hippocampus entrained by nasal respiration. J Neurosci 34:5949–5964.

Yeung M, Treit D, Dickson CT (2012) A critical test of the hippocampal theta model of anxiolytic drug action. Neuropharmacology 62:155–160.

Zhong W, Ciatipis M, Wolfenstetter T, Jessberger J, Müller C, Ponsel S, Yanovsky Y, Brankačk J, Tort ABL, Draguhn A (2017) Selective entrainment of gamma subbands by different slow network oscillations. Proc Natl Acad Sci U S A 114:4519–4524.

Zhu XO, McNaughton N (1994) The interaction of serotonin depletion with anxiolytics and antidepressants on reticular-elicited hippocampal RSA. Neuropharmacology 33:1597–1605.

